# A Single-Cell Atlas of Cell Types, States, and Other Transcriptional Patterns from Nine Regions of the Adult Mouse Brain

**DOI:** 10.1101/299081

**Authors:** Arpiar Saunders, Evan Macosko, Alec Wysoker, Melissa Goldman, Fenna Krienen, Heather de Rivera, Elizabeth Bien, Matthew Baum, Shuyu Wang, Aleks Goeva, James Nemesh, Nolan Kamitaki, Sara Brumbaugh, David Kulp, Steven A. McCarroll

## Abstract

The mammalian brain is composed of diverse, specialized cell populations, few of which we fully understand. To more systematically ascertain and learn from cellular specializations in the brain, we used Drop-seq to perform single-cell RNA sequencing of 690,000 cells sampled from nine regions of the adult mouse brain: frontal and posterior cortex (156,000 and 99,000 cells, respectively), hippocampus (113,000), thalamus (89,000), cerebellum (26,000), and all of the basal ganglia – the striatum (77,000), globus pallidus externus/nucleus basalis (66,000), entopeduncular/subthalamic nuclei (19,000), and the substantia nigra/ventral tegmental area (44,000). We developed computational approaches to distinguish biological from technical signals in single-cell data, then identified 565 transcriptionally distinct groups of cells, which we annotate and present through interactive online software we developed for visualizing and re-analyzing these data (DropViz). Comparison of cell classes and types across regions revealed features of brain organization. These included a neuronal gene-expression module for synthesizing axonal and presynaptic components; widely shared patterns in the combinatorial co-deployment of voltage-gated ion channels by diverse neuronal populations; functional distinctions among cells of the brain vasculature; and specialization of glutamatergic neurons across cortical regions to a degree not observed in other neuronal or non-neuronal populations. We describe systematic neuronal classifications for two complex, understudied regions of the basal ganglia, the globus pallidus externus and substantia nigra reticulata. In the striatum, where neuron types have been intensely researched, our data reveal a previously undescribed population of striatal spiny projection neurons (SPNs) comprising 4% of SPNs. The adult mouse brain cell atlas can serve as a reference for analyses of development, disease, and evolution.

---

Cellular specialization is central to the function of the mammalian brain. At the coarsest level, cells of different classes (for example, neurons, astrocytes, and endothelial cells) interact to maintain homeostasis and enable electrochemical communication. At the finest level, subtle specializations – such as those that distinguish neuron subtypes in the same brain region – can control behaviors such as appetite (Andermann and Lowell, 2017; Sternson, 2013), sex drive (Anderson, 2012), habit formation (O’Hare et al., 2016; Wang et al., 2011), spatial mapping (Moser et al., 2008), and associative learning (Krabbe et al., 2017; Letzkus et al., 2015). Although some of these cell populations have been characterized in detail, many more remain to be characterized or even discovered for the first time.

Systematic efforts to identify cell populations, reveal the RNA repertoires of every cell type and state, and identify markers for these populations, would help to understand the functions and interactions of cells in the brain. Comprehensive datasets would also enable new ways of probing the roles of distinct cell types in circuit function, behavior, and disease. High-throughput single-cell RNA-seq (scRNA-seq) now makes it possible to profile RNA expression in thousands of individual cells in complex tissue (Klein et al., 2015; Macosko et al., 2015; Rosenberg et al., 2018; Zheng et al., 2017). To date, single-cell gene expression studies have yielded cell-type classifications in the mouse cerebral cortex (Tasic et al., 2016a; Zeisel et al., 2015), retina (Shekhar et al., 2016), hypothalamic arcuate nucleus (Campbell et al., 2017), and amygdala (Wu et al., 2017).

Here we sought to analyze cellular diversity across a wide variety of brain regions in a uniform manner that would make it possible to learn from shared and region-specific patterns in cellular composition and gene expression. To do this, we had to overcome several challenges. First, dissociating adult mammalian brain tissue into healthy and representative cell suspensions is difficult; many scRNA-seq studies have therefore used developing mice (younger than 30 days old). However, the strong transcriptional programs associated with development can obfuscate gene-expression differences underlying the functional specializations of cell subtypes. Here we developed approaches, borrowing from techniques used for electrophysiological recordings, that allowed adult tissue to be dissociated into intact cell soma while representing all major cell classes. Second, scRNA-seq data are simultaneously shaped by real underlying cellular categories, continuously varying signals, and technical artifacts; the grouping of cells into discrete clusters often reflects unknown and non-transparent combinations of these effects. Here we developed analytical methods to dissect biological and technical influences on single-cell data sets and enable a more transparent view and revision of the resulting cellular classifications.

Here, we describe an early mouse brain cell atlas, which we created by analyzing (using Drop-seq) 690,000 individual cells from nine regions of the adult mouse brain. By comparing transcriptional patterns from within and across neuron types, we identified and validated a widely-used transcriptional state that supports axon and synapse function and discovered large-scale structure in the expression of ion channels that enable and constrain electrophysiological properties. We determined that glutamatergic neurons tend to be specialized by region in cortex, while non-neuron cell classes, such as those that make up the vasculature, can be variably specialized across cortical and subcortical areas. We also highlight the depth of individual datasets through the classification of neuron types. While we present in-depth analyses of all nine regions through DropViz, here we illustrate our insights into neuronal diversity using examples from the basal ganglia. In the globus pallidus externus (GPe) and substantia nigra reticulata (SNr), two regions where neuron types are not well-understood, we propose complete neuron-type classifications and identify selective markers for each population. In the striatum, where neuronal diversity is well charted, our data rediscovered, then built upon, accepted molecular distinctions, identifying a novel group of principal neurons that have been overlooked despite decades of research.

Our hope is that these data will advance a wide variety of efforts and nominate many unforeseen research questions. To facilitate the exploration and utilization of these data, we developed a flexible, interactive analysis platform (“DropViz” http://dropviz.org/) for comparing cell types to one another, identifying cell populations that express genes of interest, and performing many other kinds of analyses.

## RESULTS

### Isolation and Molecular Analysis of Cells for an Adult Brain Cell Atlas

To build an atlas of cell populations and cell-type-specific gene expression across many regions of the adult mouse brain, we prepared single-cell suspensions from nine brain regions (**Table S1**) and used Drop-seq (Macosko et al., 2015) to profile the RNA expression of 690,207 individual cells (**Figure 1A,B**). We profiled cells from the Frontal and Posterior Cerebral Cortex (FC and PC), Hippocampus (HP), Thalamus (TH), Cerebellum (CB), and the Basal Ganglia (consisting of the Striatum [STR], Globus Pallidus Externus / Nucleus Basalis [GP/NB], Entopeduncular Nucleus / Subthalamic Nucleus [EP/STN] (Wallace et al., 2017), and Substantia Nigra/Ventral Tegmental Area [SN/VTA]) (**Figure S1**).

**Figure 1.**
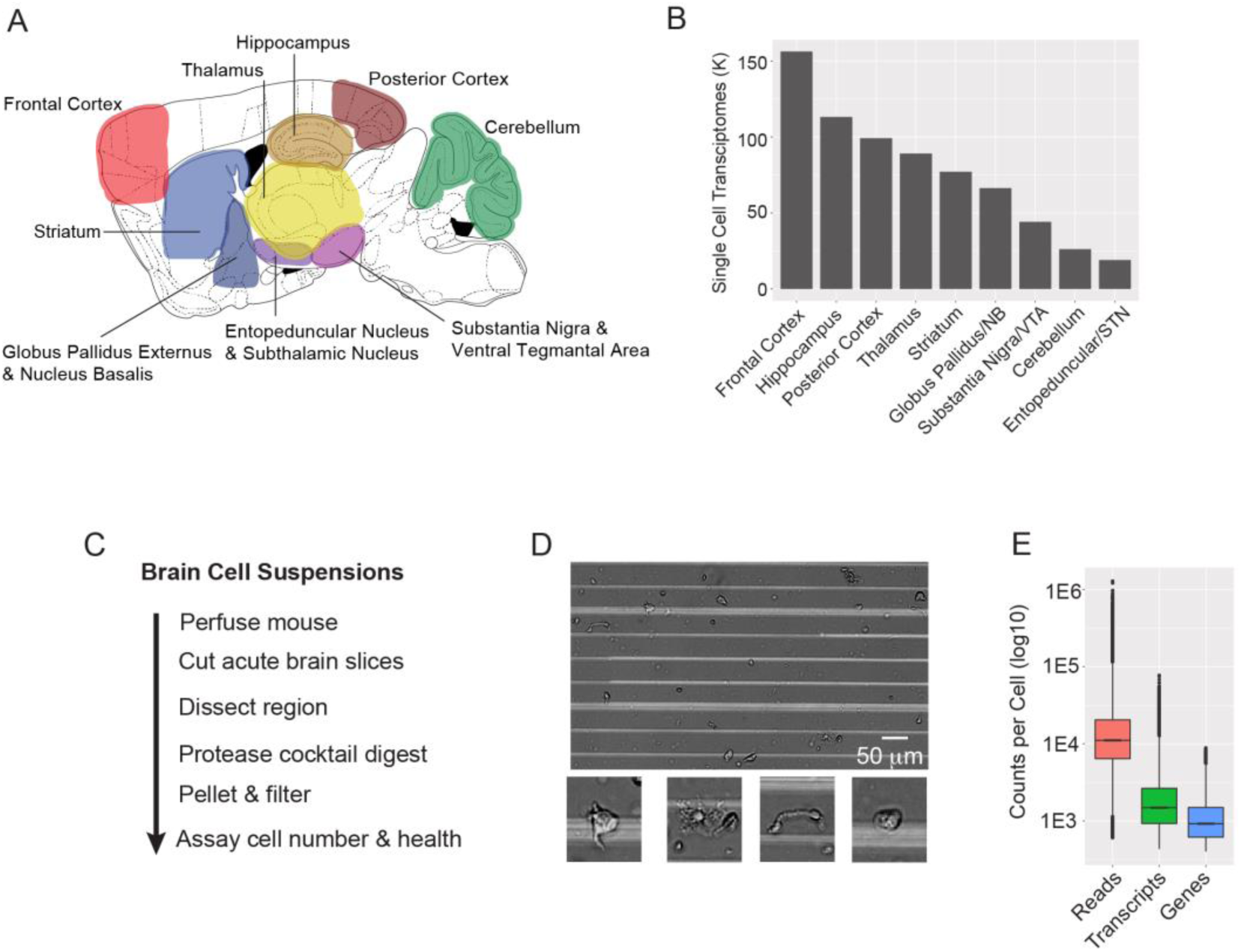
Single-cell transcriptional profiling of the adult mouse brain using Drop-seq. A. Sagittal brain schematic illustrating the n=9 profiled regions. Detailed schematics are presented in **Figure S1**.
B. Plot illustrating the number of single-cell transcriptomes per region.
C. Flowchart describing how acute single-cell suspensions are prepared from adult mouse brain.
D. Example single-cell suspension generated from frontal cortex that contained little debris and diverse cellular morphologies. Bottom, enlargements of example cells.
E. Distribution of sequencing reads, transcripts, and genes for all cells (n=690,207)

We generated cell suspensions from adult male mice (60–70 days old; C57Blk6/N) by adapting protocols from single-cell patch-clamp recording (Carter and Bean, 2009). We optimized the digestion time for each region to maximize cell recovery rates (see **Methods**, **Figure 1C**). Tissue was prepared in a buffer lacking Ca^2+^ with ionic concentrations designed to avoid activity-induced toxicity by maintaining voltage-gated Na channels in an inactivated state [Estimated V_m_ = −30.5 mV, **Methods**]). The resulting cell suspensions, which recovered intact 40–50% of cells from most tissues (cortex: 0.46 ± 19 mean ± sem; striatum: 0.39 ± 20, **Figure S4A-C**), had cells with the characteristic morphologies of neurons, astrocytes, and oligodendrocytes, and were largely free of debris (**Figure 1D**). We prepared 3-7 replicate suspensions of each brain region (44 total).

We performed Drop-seq analysis of 690,207 individual cells from these 44 cell suspensions (Macosko et al., 2015). We generated and analyzed 13 billion sequencing reads from the resulting Drop-seq libraries, ascertaining 1.45 billion distinct mRNA transcripts (UMIs), which arose from 31,767 distinct genes. We ascertained an average of 17,480 reads (median = 10,824), 2,218 mRNA transcripts (median= 1,450 UMIs), and 1,169 genes per cell (median = 900) (**Figure 1E**).

### Cell-class Composition of Nine Adult Brain Regions

We first separately analyzed data for each of the nine brain regions, analyzing each region in two stages (here termed “global clustering” and “subclustering”). The first round of analysis, which is relatively straightforward (**Methods**), grouped cells into 8-11 broad classes that were then easily identified by known marker genes; these broad classes included neurons, astrocytes, microglia/macrophages, oligodendrocytes, polydendrocytes (oligodendrocyte progenitor cells), and components of the vasculature – endothelial cells, fibroblast-like cells, and mural cells (Abbott et al., 2006; Marques et al., 2016; Vanlandewijck et al., 2018; Wälchli et al., 2015). For example, the hippocampus yielded cells from all [11 cell classes](http://dropviz.org/?_state_id_=2b493beb4e780071) (**Figure 2A-C**), including three local cell classes, of which two are native to the ventricle – the choroid plexus and ependymal cells, which had been sampled from ventricular tissue proximal to the hippocampal dissections, and an additional cell class undergoing adult neurogenesis (Habib et al., 2016; Hochgerner et al., 2018; Ming and Song, 2011). Assignment of cells to these broad classes was robust to analysis parameters (**Figure S2**).

**Figure 2.**
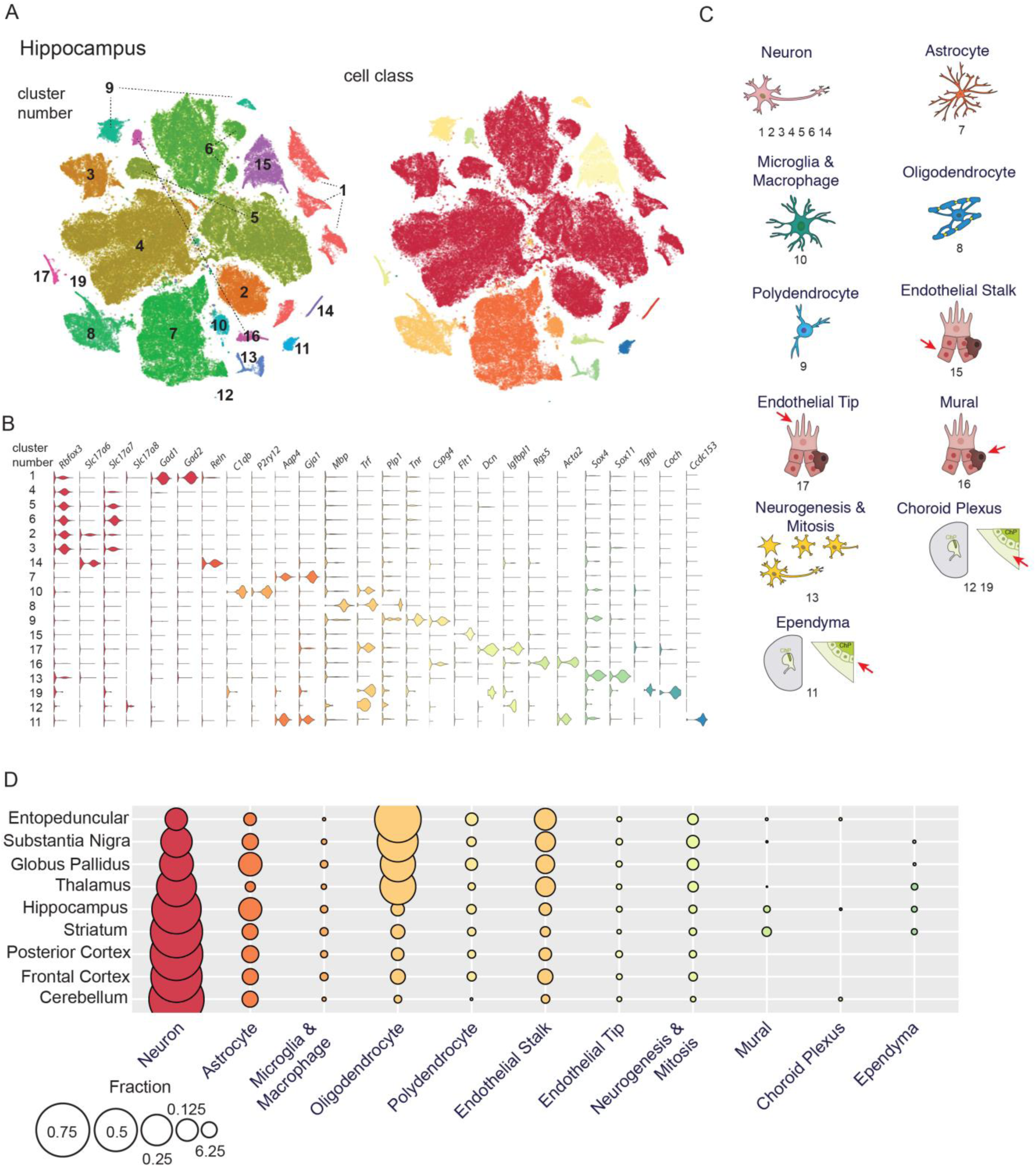
Comprehensive identification of brain cell classes. A. T-distributed stochastic neighbor embedding (tSNE) plot of gene expression relationships amongst the n=113,171 cell hippocampus dataset based on the first round of ICA-based clustering. The resulting "global" clusters (colored independently and numbered 1-19) are each associated with major cell classes of the brain. Right, global clusters color-coded by cell class.
B. Violin plots showing example gene markers that distinguish across and within brain cell classes (log10). Genes are color-coded by cell class. Global clusters are ordered by cell class.
C. Cartoons of the major cell classes of the brain. Numbers below indicate the corresponding global clusters from hippocampus.
D. Dot plots displaying the proportional representation of individual cell classes across regions.

Distinct brain regions yielded the major cell classes in different proportions. (**Figure 2D**). The proportion of neurons varied inversely with those of oligodendrocytes, endothelial cells, and mural cells (**Figure 2D**); a natural explanation for this is the regional variation in the abundance of grey matter, with oligodendrocytes (as expected), endothelial cells, and mural cells being more abundant in white matter than in grey matter. Fibroblast-like cells comprised a similar proportion of all cells in each region. Polydendrocyte and astrocyte abundance appeared to vary independently of other cell classes, exhibiting enrichment in the GPe as expected based on earlier findings (Cui et al., 2016; Lange et al., 1976). Small fractions of choroid plexus and ependymal cells were sampled from ventricle-adjacent regions, while sparse neurogenic populations were observed in regions adjacent to the subventricular and subgranular zones (frontal cortex, striatum, and hippocampus) (Ming and Song, 2011).

### Inference of Cell Types and States Using Independent Components Analysis

The systematic recognition of more-subtle (yet consistent) patterns of variation among cells of the same class presents formidable analytical challenges – particularly when these distinctions are to be made in systematic, inductive (unsupervised), data-driven ways (Mayer et al., 2015; Satija et al., 2015; Shekhar et al., 2016; Tanay and Regev, 2017; Tasic et al., 2016b). The size and diversity of the data sets for these nine brain regions, and the generation of many biological replicates for each region, made apparent many limitations in diverse analysis methods. We initially found clusters that were specific to experimental replicates or that were defined by transcriptional signatures that we recognized as artifacts of tissue digestion (see below). We sought an analysis strategy that would make it possible to (i) dissect biological from technical contributions to the expression data, so that technical contributions could be excluded from subclustering analysis; and (ii) generate transparent and meaningful intermediate data sets (downstream of input expression data, and upstream of clustering outcomes) that could be critically evaluated – and, ideally, be meaningful objects of analysis in their own right.

We therefore developed an analysis strategy based on independent components analysis (ICA)(**Figure 3A,B**). ICA reduces large datasets to a smaller number of dimensions in which entities (here cells) have score distributions that are maximally structured – as measured by deviation from a normal distribution (generally due to a spikey or clustered distribution of the cells in that dimension) – and statistically independent (Hyvärinen, 1999). Each of the inferred independent components (ICs) is a weighted combination of many genes (the weight of each gene’s contribution to an IC is the gene “loading”). Each cell’s gene-expression profile is reduced to a weighted sum of these ICs, with each IC having a score for each cell that reflects the expression of that combination of genes in that cell.

**Figure 3.**
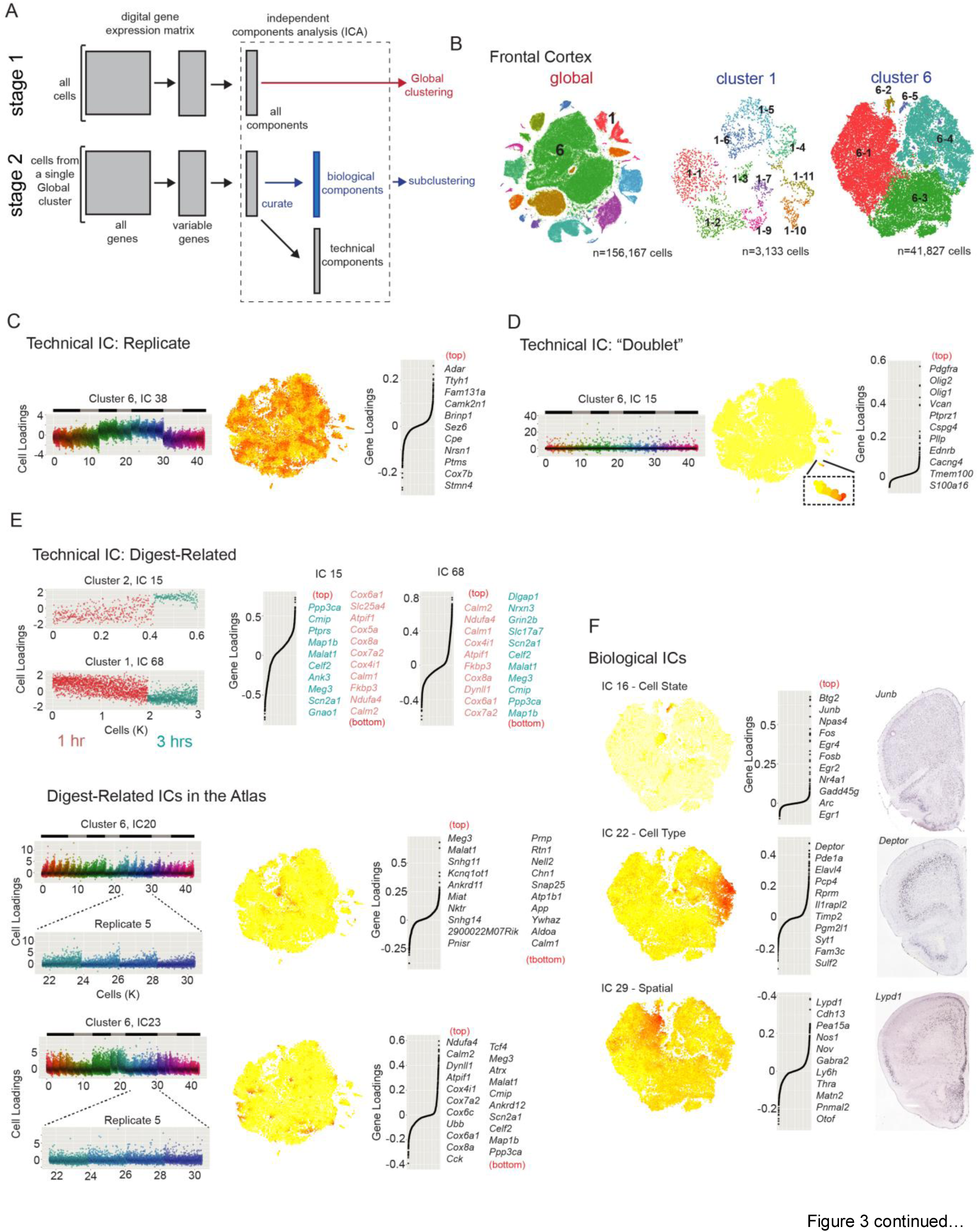
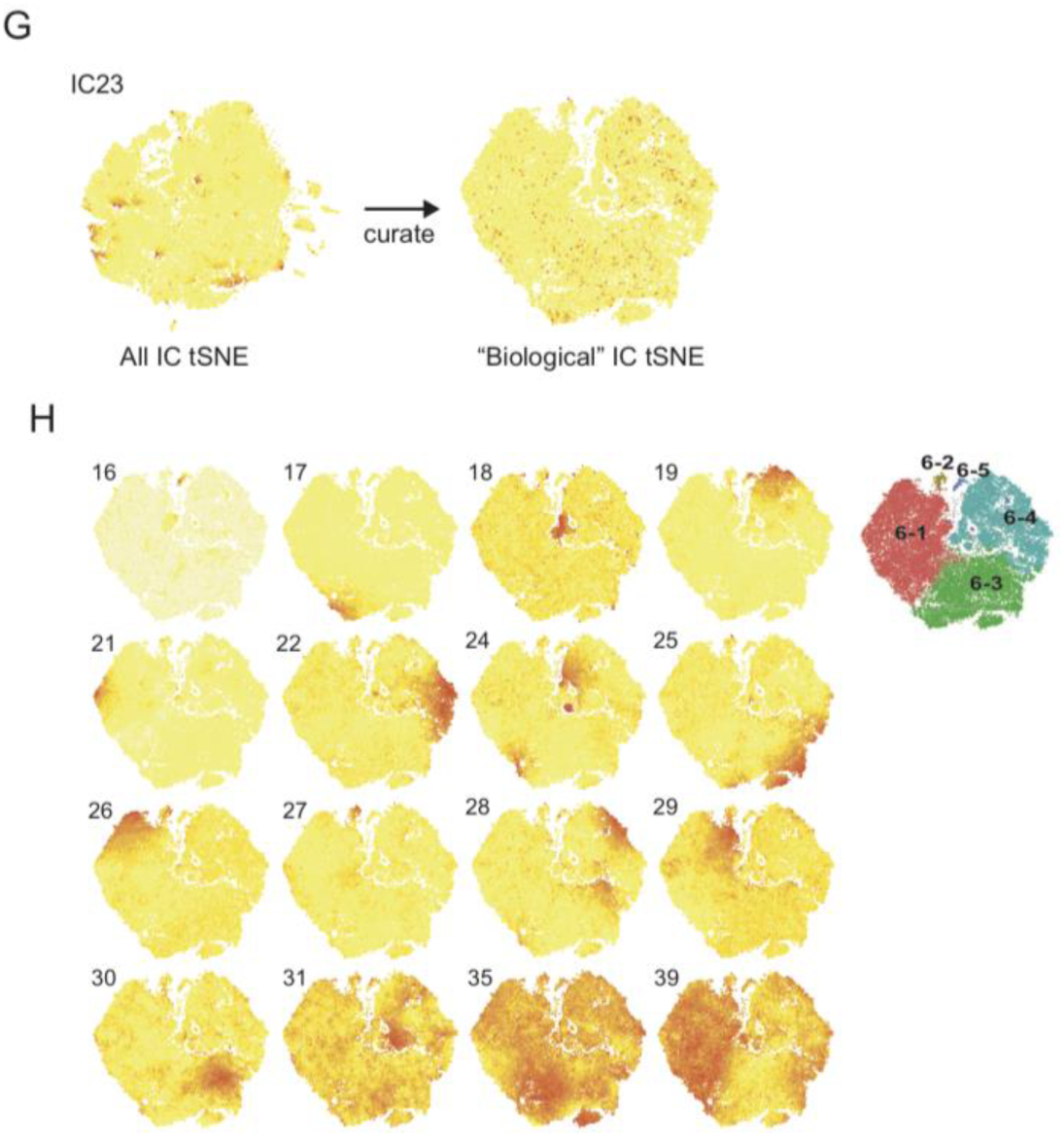
Independent component analysis identifies transcriptional programs allowing transparent description of within-cell class molecular diversity. A. Workflow for semi-supervised two-stage ICA-based signal extraction and clustering. In the first stage (“global clustering”, red arrows), the full digital gene expression (DGE) is clustered into “global” clusters using a reduced space generated by ICA (Methods). These “global” clusters represent cell classes. The process is then repeated independently for each of the global clusters with >200 cells (“subclustering”, blue arrows), except that the resulting Independent Components (ICs) are manually curated into two categories: “technical” components representing signals associated with the 1) replicate (batch effects); 2) libraries containing two cells (“cell-cell doublets”) or single-cells with unique transcriptional signatures (“outliers”) or 3) the preparation process for acute tissue. “Biological” signals originate from cell types, cell states, or from spatial anatomical relationships. Only “Biological” ICs are used as input to subclustering. Subclustering is performed over a range of neighborhood sizes and resolutions, generating a portfolio of potential solutions. The single solution with the clearest correspondence between Biological IC cell loading and subclusters is selected (Figure S3).
B. Color-coded tSNE plots for frontal cortex global clustering (left) and two representative subclusterings, GABAergic interneurons (middle, cluster 1) and glutamatergic layer 2/3 and some layer 5 neurons (right, cluster 6). (C-E) Examples of “technical” ICs from frontal cortex.
C. Example of an independent component (IC 38, cluster 6) representing a replicate-based signal. Left, plot of cell-loading scores (y-axis). On the x-axis, cells are ordered by library size (largest to smallest) within each sequencing pool (color-coded). Sequencing pools from the same mouse are grouped with black and gray bars above. Middle, cell-loading scores plotted on the subcluster 6 tSNE (tSNE constructed from all ICs). Higher loadings are shown in darker red. Right, gene-loading plot. The n=10 genes with the highest scores are shown at right.
D. Example of a component (IC 15, cluster 6) representing a “doublet” signal. Plots and layout are similar to C. IC 15 loads heavily on a small number of cells across mice and sequencing pools. The top loading genes are markers of the polydendrocyte cell class, suggesting a signal that represents a “doublet” transcriptome consisting of layer 2/3 neurons and polydendrocytes. Inset shows area of the tSNE plot with high IC 15 loading.
E. Experimental identification of protease digest-related technical ICs. Top, frontal cortex tissue was under- (1 hr) or over- (3 hr) digested before making single-cell suspensions and Drop-seq libraries. The datasets were co-analyzed. Subcluster IC loadings that correlate with digest time were identified (Figure S3C-E). Example digest-related ICs (IC 15, cluster 2, Syt6+ deep-layer glutamatergic) and (IC 68, cluster 1, Nptxr+ upper-layer glutamatergic) show contributions from similar genes. Genes related to ATP synthesis and Calmodulins (eg., Cox6a1, Ndufa4, Calm2) load on the under-digested cells, while another set of genes including the nuclear-enriched Meg3 and Malat1 load highly onto the over-digested condition. Bottom, similar digest-related ICs were commonly observed in atlas subclustering analyses, suggesting heterogeneity in cellular transcriptional response to identical preparation conditions. Two examples of digest-related ICs from frontal cortex cluster 6 (20 and 23). Left, cell-loading plots and insets highlight the correlation of IC loading to library size. IC 20 tends to load on smallest libraries, while IC 23 loads on the largest. Middle, cell-loadings for IC 20 and 23 demonstrate that digest-related signals contribute to local relationships within the subcluster 6 tSNE. Right, gene-loading plots for the 10 top and bottom loading genes indicate a similar signal to that identified in over and under-digested experiments.
F. Examples of heterogeneous “Biological” ICs from frontal cortex cluster 6, representing a cell state (top, IC 16), cell type (middle, IC 22), and spatial anatomical signal (bottom, IC 29). For each example, a cell-loading tSNE plot, gene loading plot, and in situ hybridization experiment for a top-loading gene are shown from left to right. IC 16 corresponds to the immediate early gene (IEG) signal. The IC 22 signal originates from layer 5a glutamatergic neurons, exemplified by the spatially restricted expression of top-loading Deptor. IC 29 represents a spatial signal. Top-loading genes such as Lypd1 exhibit an expression gradient from medial (highest) to lateral (lowest). ISH data from the Allen Institute Mouse Atlas.
G. Removing technical ICs prevents spurious transcriptional similarities. Example tSNE cell-loading plot for cluster 6 digest-related IC 23 before and after IC curation. Before curation, cells with high loading of digest-related IC 23 (E, bottom left) are locally grouped. After curation, technical ICs (including IC 23) are removed, creating a different tSNE plot and preventing the digest-related effects from contributing to clustering.
H. Correspondence between heterogeneous transcriptional signals (Biological ICs) and subclusters identified by modularity-based clustering (Methods). tSNE cell loading plots for each of the n=16 Biological ICs from frontal cortex cluster 6. Top right, the resulting plot in which the n=5 subclusters are identified. The portfolio of alternative subcluster solutions is shown in Figure S3G.

We found that many individual ICs corresponded to easily recognized biological phenomena (**Figure 3C-F**)(Adamson et al., 2016). For example, among cells from frontal cortex [cluster 6](http://dropviz.org/?_state_id_=ca88a79461e5cc9f) (glutamatergic neurons from layer 2/3 and 5a), we identified ICs whose strongly loading genes were readily recognized as known markers for specific [cell types](http://dropviz.org/?_state_id_=4e03a8081aac89ef), [cell states](http://dropviz.org/?_state_id_=2e3edcfb5f07fc75) or [spatial gradients](http://dropviz.org/?_state_id_=3c5919ca5d557170) across anatomical axes (**Figure 3F**). Other ICs exhibited cell-loading distributions that immediately suggested technical effects, such as ICs that distinguished 1) cells (of all classes) from different replicate preparations from the same brain region; 2) single-cell RNA libraries of different sizes; 3) experimentally-identified effects of tissue preparation (**Figure S3C-E**); or 4) cell-cell “doublets” (**Figure 3C-E**). (By contrast, in principal components analysis (PCA), which optimizes components to maximize variance without regard to statistical independence or data distribution, distinct biological signals are typically distributed across many components and mixed with technical sources of noise (**Figure S3A**); PCs still carry maximal information forward to downstream clustering algorithms while reducing data dimensionality, but are not intended to parse statistically distinct signals into separate components). We found that the interpretability of individual ICs allowed us to distinguish ICs corresponding to presumed endogenous signals (which we call here “biological ICs”) from ICs related to technical signals described above. Moreover, by removing contributions from artifact ICs, we reduced spurious noise (**Figure 3G**).

We analyzed the data from each class and brain region (109 analyses total, **Table S9**) using a semi-supervised ICA, in which we excluded 1,157 ICs as recognizably technical in origin in that they corresponded doublets of cells from different classes (n=759), individual outlier cells (n=99), or replicate-specific or tissue preparation artifacts (n= 299) (**Table S11**). We classified the remaining 601 ICs as “biological ICs”, though it remains possible that additional technical influences on the data were not recognized during curation. We then clustered the cells for each of the 109 brain-region-specific, cell-class-specific analyses based on their combination of loadings for the “biological ICs” in the respective analysis.

To group cells into subclusters, we used network-based clustering (Waltman and van Eck, 2013), which has been utilized in earlier studies (Shekhar et al., 2016). For each subcluster analysis, we used only the putative “biological ICs” as dimensions for graph construction. To create a transparent relationship between biological ICs and subclusters, we utilized clustering-resolution parameters that maximized one-to-one relationships between subcluster assignments and underlying ICs (**Figure 3H** and **Figures S3E,F**). The 109 separate subclustering analyses of the cell classes from each of the nine regions identified 565 total subclusters **(Figure 4A** and **Table S10**). Of these subclusters, [323 were neuronal](http://dropviz.org/?_state_id_=633342e29c7798c2) and were derived from 368 biological ICs (**Figure S4**). Hierarchical clustering of the expression profiles for subclusters (aggregated across their constituent cells) demonstrated that clustering is driven primarily by cell class and not by brain region (**Figure 4B**).

**Figure 4.**
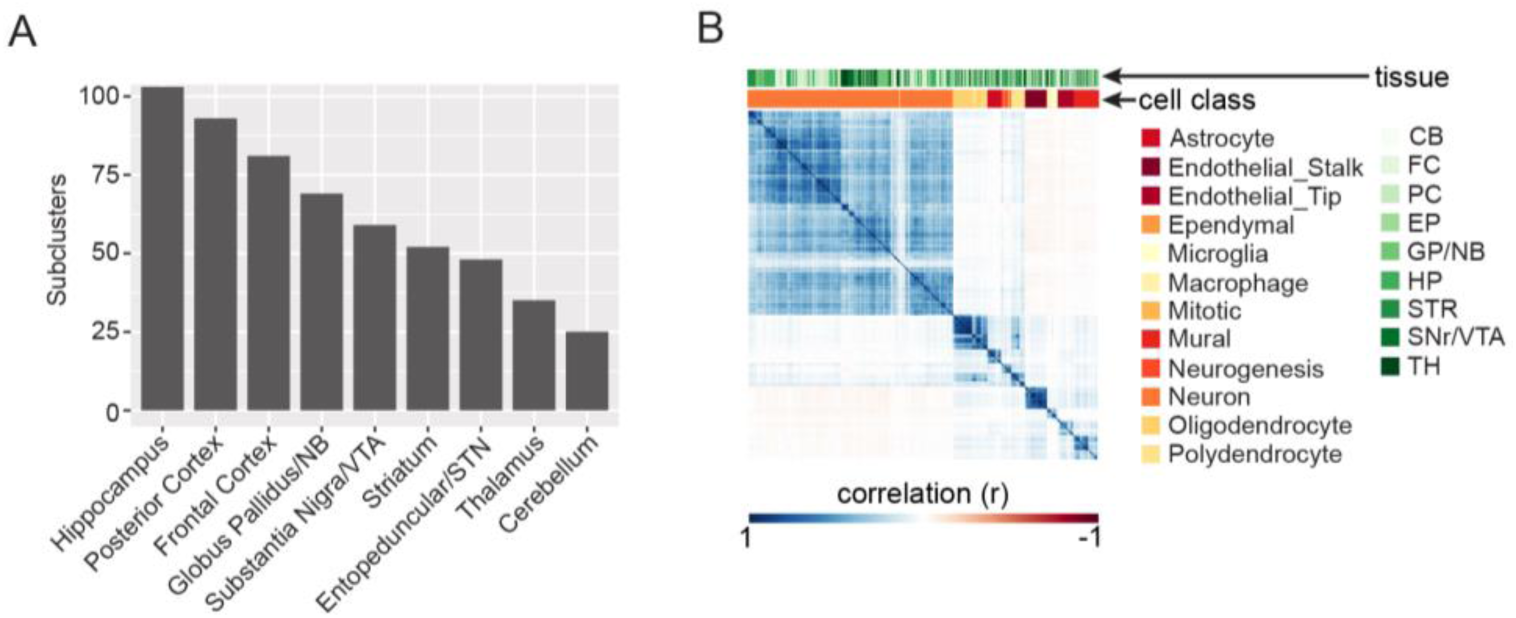
Transcriptional diversity by region and cell class. A. Number of subclusters by region.
B. Transcriptional correlations across atlas subclusters are largely explained by cell class and not region of origin. Hierarchical clustering diagram showing pairwise Pearson correlation scores calculated pairwise between average atlas subclusters. The analysis was restricted to genes with significantly variable expression (**Methods**). Color-coded bars at the top of the plot display the ordered region/cell class assignments for the subcluster.

### Characteristics of the Cells of the Blood-Brain Barrier

Non-neuronal cells, including glia and endothelial cells, exhibited broadly consistent expression signals across brain regions. To better appreciate diversity among non-neuronal classes, we grouped single-cell libraries from each region by cell class and performed semi-supervised ICA on each of the seven datasets independently, identifying a total of 53 biological ICs (**Figure 5A** and **Figure S5A-F**). We focus here on cell classes that form the blood-brain barrier – mural, fibroblast-like (aka vascular leptomeningeal cells (Marques et al., 2016)) and endothelial cells – because they are diseaserelevant (Sweeney et al., 2016) and remain incompletely characterized (**Figure 5B**).

**Figure 5.**
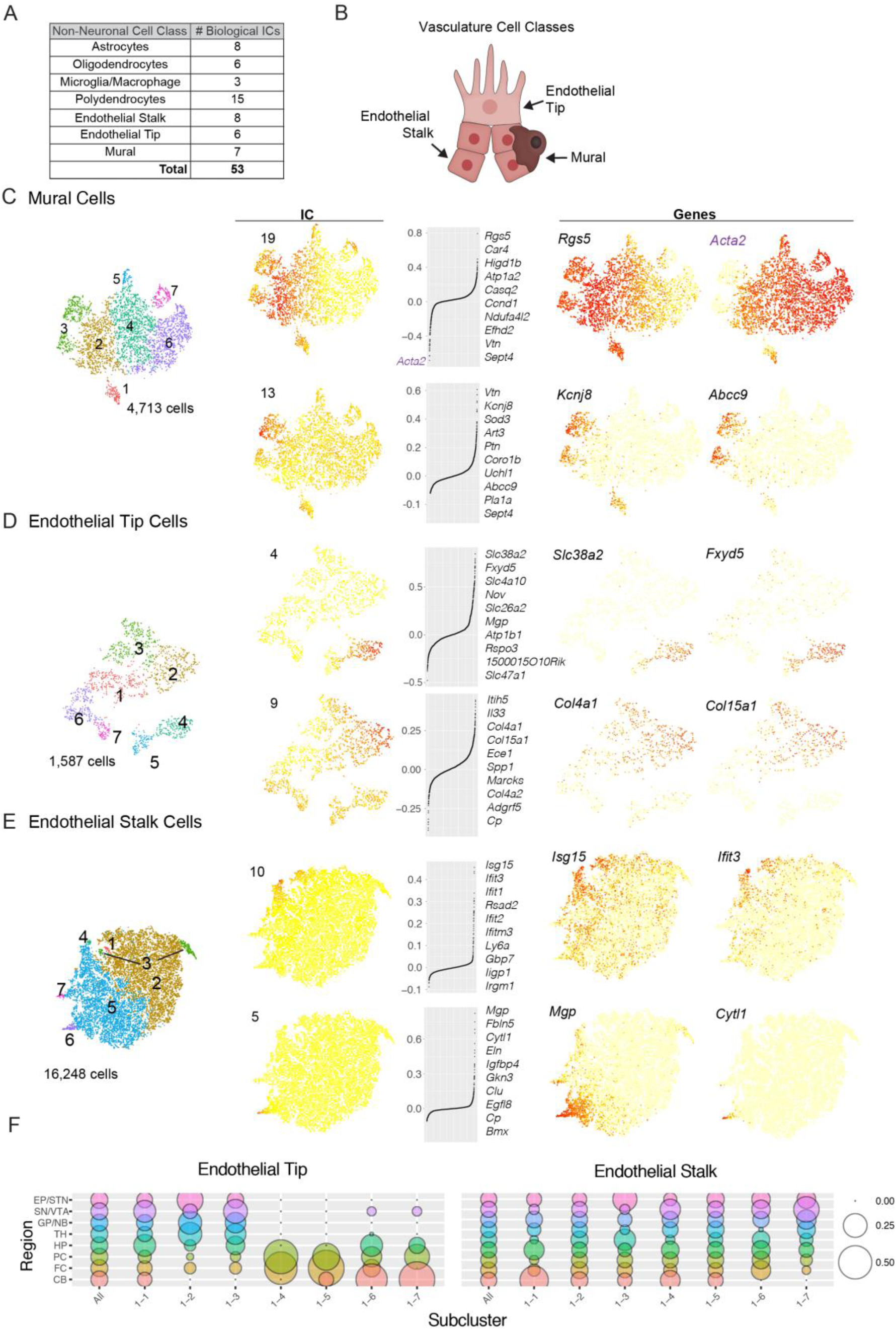
Comprehensive description of transcriptional diversity within non-neurons as illustrated by cell classes of the vasculature. (A) Table showing the number of biological ICs curated for each non-neuronal cell class. A description of non-neuronal ICs is provided in **Figure S5**. (B) Cartoon of vasculature cell classes. (C-E) Subcluster assignments and examples of two biological ICs for each vasculature cell class. Subclusters (color-coded), IC cell-loadings, and gene expression values displayed on tSNE plots. Left, subcluster assignments. Middle, IC cell- and gene-loadings. For each IC, the top ten loading genes are listed. Right, expression plots for individual genes. For Mural Cell IC 19, the bottom loading gene Acta2 is shown in purple. (F) Dot plots illustrating fractional representation of cells from each region contributing to each particular fibroblast-like and endothelial subcluster. Similar plots for other non-neuronal cell classes are shown Figure S5H.

Mural cells are morphologically diverse cells embedded within the basement membrane of the endothelium. Mural cells control vascular development, stability, and homeostasis, and exhibit pathology in many disease states (Sweeney et al., 2016; Trost et al., 2016). We identified 7 mural cell subclusters from 7 biological ICs (n=4,713 cells, **Figure 5C and Figure S5E**). Mural cells are known to have two subtypes: pericytes, which associate with small-diameter capillaries and are enriched for *Rgs5* expression, and smooth muscle alpha actin (SMA) cells, which associate with larger-bore vasculature and express *Acta2* (Bondjers, 2006; He et al., 2016; Hughes and Chan-Ling, 2004; Nehls and Drenckhahn, 1991). Within arteries and veins, *Acta2* expression is graded, with highest expression in large-bore arteries and lowest expression in veins (Vanlandewijck et al., 2018). One independent component (IC 19) appeared to correspond to the arterial versus veinous distinction within SMA cells, since *Rgs5* and *Acta2* were the most strongly weighted genes (**Figure 5C**). Notably, the data suggested that this distinction may be continuous rather than categorical, as the expression levels of *Rgs5* and *Acta2* were graded and the cell scores for this IC were continuously rather than bimodally distributed across these cells (**Figure 5C**)(Trost et al., 2013; Vanlandewijck et al., 2018). (For those downstream analyses that require comparisons of discrete cell populations, we discretized this signal into three subclusters (2,4,6), but “clusters” cannot be reflexively equated with “discrete cell types”.) A second component (IC 13) appeared to correspond to the pericyte versus SMA distinction, since strongly weighted genes included known pericyte markers such as *Vtn* (**Figure 5C**)(Vanlandewijck et al., 2018). Presumed pericytes (cluster 3) expressed the pericyte marker *Rgs5* and lacked expression of the SMA marker *Acta2*. Other enriched genes are suggestive of specialized function. For example, expression of genes encoding a potassium channel potently activated by intracellular diphosphate levels (*Kcnj8* and *Abcc9*) and an ADP-ribosyltransferase (*Art3*) indicate this pericyte population has signaling machinery that couples dinucleotide metabolites to membrane potential and post-translational modification. Interestingly, the expression of *Rgs5* and *Acta2* was not always inversely correlated: a subpopulation of mural cells (cluster 1) expressed both *Rgs5* and *Acta2*, and also uniquely expressed genes such as *Aldh1a1* (**Figure S5E**), suggesting additional diversity within mural cells with no known function or anatomical correlate (Vanlandewijck et al., 2018).

While endothelial cells have been long-known as the constituent cell class of blood vessels, fibroblast-like cells are a recently-described population with unknown function that inhabit the perivascular space of brain arteries and veins but not capillaries (Vanlandewijck et al., 2018). We found 7 subclusters each of endothelial (n=16,248) and fibroblast-like (n=1,587) cells (**Figure 5D,E**). Among the fibroblast-like cells, two clusters (4 and 5) were notable for their expression of genes encoding a variety of membrane transporters (e.g., *Slc38a2*, *Slc4a10*, *Slc26a2*, and *Slc47a1*) and pumps (e.g., *Fxyd5* and *Atp1b1*) (**Figure 5D**). To varying extents, clusters 1, 2, and 3 expressed genes involved with extracellular matrix (ECM) secretion, angiogenesis, and contraction (**Figure S5F**), such as the basement membrane collagen genes (*Col4a1*, *Col4a2*, and *Col15a1*) (**Figure 5D**). Interestingly, cluster 3 expressed a distinct set of collagen genes (*Col1a1* and *Col3a1*) (**Figure S5F**). These examples suggest that subsets of fibroblast-like cells are transcriptionally specialized to affect membrane transport and ECM production; ECM secretion may involve coordinated expression of distinct sets of collagen genes.

A recent scRNA-seq analysis by Vanlandewijck et al. suggested a continuous transcriptional relationship between endothelial cells (**Figure 5E**) associated with arteries, capillaries and veins (Vanlandewijck et al., 2018). We identified independent components that exhibit strong contributions from genes with expression restricted to arterial (IC 5 and 20) and veinous endothelial populations (IC 3 and 12) (Vanlandewijck et al., 2018), suggesting transcriptional heterogeneity related to blood vessel type (**Figure S5G**). Genes contributing to IC 20 (e.g. *Tm4sf1* and *Slc38a5*) exhibited continuous, reciprocal expression levels across endothelial cells, indicating a signal that likely encodes a smooth molecular transition across blood vessel types (**Figure S5G**). Other ICs had strong contributions from genes with more bimodal levels of expression across subsets of endothelial cells. For example, one subpopulation (cluster 6) expressed the highest levels of the artery marker *Bmx* and exclusive expression of *Cytl1* (**Figure 5E**). Other enriched genes include those that are implicated in growth-factor dependent vascular remodeling (*Mgp*, *Fbln5*, *Eln*, *Igfbp4*, and *Clu*) (**Figure S5G**)(Boström et al., 2004; Contois et al., 2012; Fitch et al., 2004; Fu et al., 2013; Guadall et al., 2011; Karnik et al.; Vanlandewijck et al., 2018), suggesting that this arterial subpopulation may be specialized to coordinate a component of angiogenesis. We also observed endothelial subpopulations that contained cells expressing different blood vessel markers and thus could represent a transcriptional specialization shared across vessel types. For example, one subpopulation expressed genes related to host immunity; these included genes encoding interferons (*Ifit3*, *Ifit1*, *Ifit2*, and *Ifitm3*), GTPases induced by interferons (*Ligp1*, *Irmg2*, and *Gbp7*), and other proteins involved in the anti-viral response (*Isg15* and *Rsad2*) (**Figure 5E**). Cluster 4 had the highest levels of expression of these genes, including *Isg15* and *Ifitm3.* These examples define transcriptional specializations within endothelial cells related to blood vessel type, processes within particular vessel types (angiogenesis) and processes that span vessel types (host defense). Other groups might reflect cell states specialized for iron handling, calcium signaling, and the stress response (**Figure S5G**).

Functional specializations within endothelial, glial, and other non-neuronal cell classes could be a ubiquitous feature of the adult mouse brain, or could be enriched in particular brain regions. We compared the abundance of fibroblast-like and endothelial cells from each brain region within each of the clusters (**Figure 5H**) (results for astrocytes, oligodendrocytes, polydendrocytes, microglia/macrophages and mural cells are in **Figure S5H**). While endothelial cell subpopulations appeared in similar relative abundance across regions, fibroblast-like cell subpopulations exhibited different contributions from cortical and subcortical areas: for example, the cortex and hippocampus contributed disproportionately to the fibroblast-like population that more strongly expressed genes with membrane-transport functions (cluster 6), while collagen-expressing cell populations (clusters 2 and 3) came largely from the basal ganglia and thalamus (**Figure 5H**).

### A Neuronal Transcriptional Program Related to Axon Function

Cell states involve constellations of genes that are transiently co-expressed to enact cellular functions.

In neurons, the most well-studied cell state, involving the expression of the immediate early genes (IEGs), is activated by Ca^2+^ influx after neuronal action potential firing (Bading, 2013; Hrvatin et al., 2017). IEGs (such as *Fos* and *Junb*) encode transcription factors that orchestrate a cellular response to neuronal activity. IEG expression in response to action potential firing is known to be largely uniform across neuronal types(Hrvatin et al., 2017) and brain regions and exhibits little background expression, making it straightforward to detect in the Drop-seq data. Though IEG induction has been reported as an artifact of cell dissociation(Hrvatin et al., 2017; Wu et al., 2017), we had sought to avoid promiscuous expression by digesting tissue in a buffer which lacked Ca^2+^ and inactivated channels necessary for action potential firing (**Methods**). Among the cells in this analysis, elevated expression of IEGs was observed in only a small fraction of the neurons of each type. For example, in [FC layer 2/3 and 5a](http://dropviz.org/?_state_id_=3a1965189e5cc594) (cluster 6, **Figure 3**) or [HP CA3](http://dropviz.org/?_state_id_=780f658ffa283012) (cluster 6, **Figure S4C**) – regions prone to epileptiform activity due to recurrent, excitatory connections – IEG^+^ neurons comprised less than 1.0% of the total population.

Neurons could in principle have other common transcriptional dynamics or cell states. To identify such patterns, we looked for broad transcriptional patterns that, despite being ascertained in many different brain regions or neuronal classes, involve similar combinations of genes. More specifically, we calculated pairwise IC-IC correlations of gene loadings for 368 biological ICs ascertained from neuronal cell classes (**Figure 6A**). One block of 15 highly inter-correlated ICs clearly represented the IEG signal in different regions (a positive control). Two additional correlation blocks were prominent. One of these arose entirely from thalamic neurons (of diverse types), suggesting a tissue-specific transcriptional program (that we did not further analyze). We focused on the other correlation block, which consisted of ICs from many brain regions and neuronal types.

**Figure 6.**
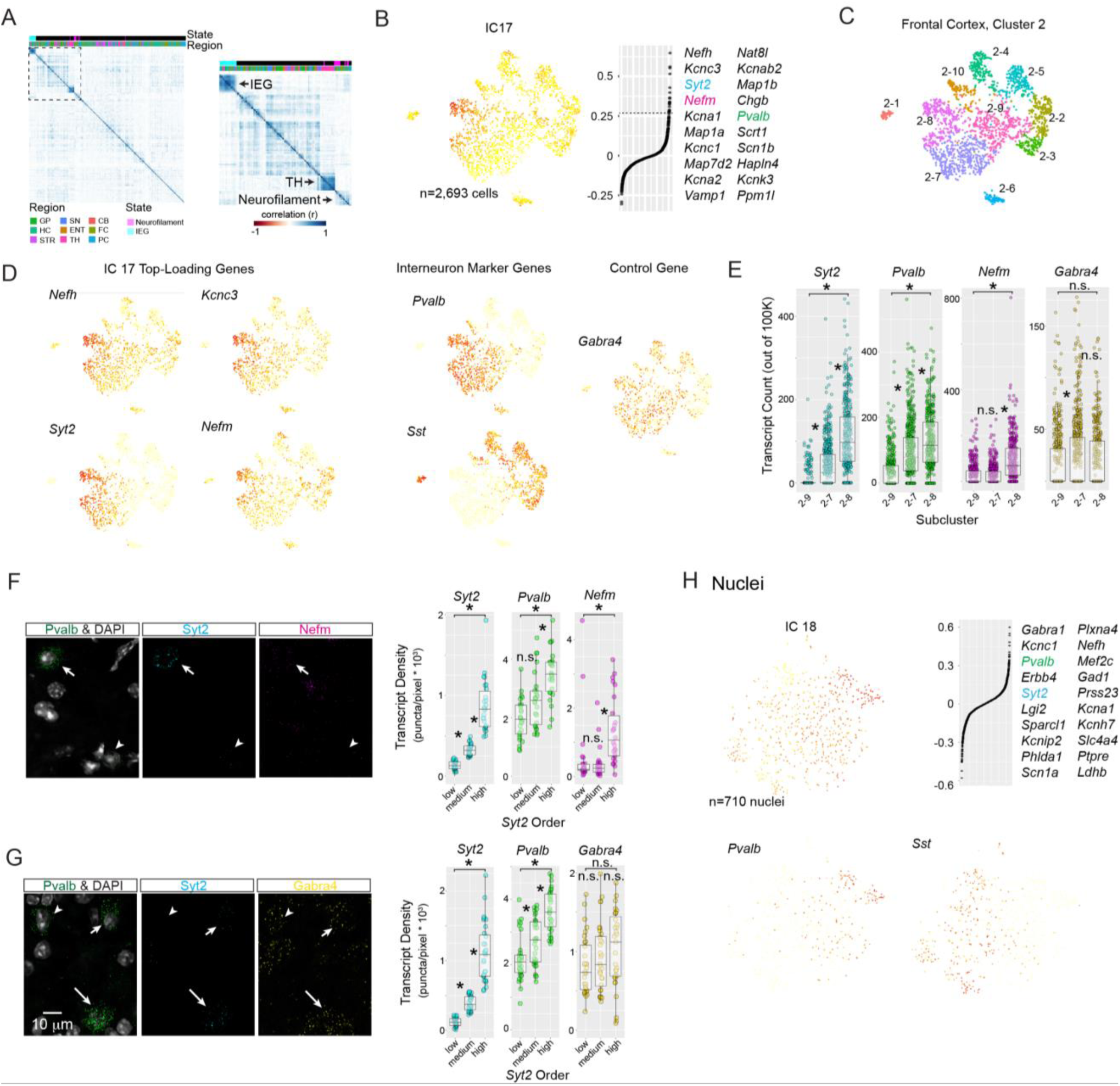
A prevalent expression state in neurons related to axon structure and presynaptic function. (A) Identification of a novel transcriptional state in neurons using biological ICs. ICs encoding transcriptional states utilized by different types of neurons can be identified by high correlations in expressed genes (gene-loadings). Hierarchical clustering diagram displaying pairwise Pearson correlations of gene-loading scores for all biological ICs curated from neuronal subclustering (n=45 subclustering analyses, n=323 total subclusters). Right, enlargement of the boxed region. IC gene-loading correlation blocks identify the immediate early gene (“IEG”) transcriptional state, with IC contributions from different regions. Another correlation block (“TH”) identifies a thalamus-specific collection of ICs. A third correlation block (“Neurofilament”) has IC contributions from different regions and top-loading genes containing neurofilament subunits and other proteins involved in Ca2+ handling, vesicle exocytosis, and membrane excitability. ICs were color-coded for states by assessing the presence of state-marker genes in top loading genes. (B) The Neurofilament transcriptional signal (IC 17) in frontal cortex Sst+/Pvalb+ interneurons (Cluster 2). Left, IC 17 cell-loadings displayed on subcluster tSNE plot. Right, gene-loading plot, with the top 20 loading genes shown at right. Syt2, Nefm, and Pvalb are color-coded to match ISH experiments below. (C) Color-coded subcluster identities for frontal cortex cluster 2. N=10 subclusters were based on n=9 biological ICs. The graded loading of IC 17 is discretized into subclusters 2-8, 2-7, and 2-9. (D) Expression plots for single genes. Left, n=4 top loading genes for IC 17 exhibit gradients of expression in Pvalb+ but not Sst+ interneurons (middle). Right, Gabra4 – a control gene that does not load onto IC 17 – does not exhibit expression gradients across Pvalb+ subclusters. (E) Quantitative comparison of Neurofilament (Syt2, Pvalb, and Nefm) and control gene (Gabra4) single-cell transcript counts across Pvalb+ subclusters from Drop-seq. Transcript means were compared with a two-way Anova followed by a Tukey Honest Significance Difference Test to calculate a p-value for differences across means. Asterisk, P < 0.05; n.s P > 0.05. (F-G) Neurofilament gene and control gene in situ transcript count experiments within Pvalb+ frontal cortex cells using smFISH. Left, example single planes from confocal stacks. Right, quantification of transcript densities. Pvalb+ cells were split into n=3 groups based on Syt2 levels (low, medium, and high) mimicking subclusters 2-9, 2-7, and 2-8 from the atlas. Transcript densities were compared with a two-way Anova followed by a Tukey Honest Significance Difference Test to calculate a p-value for differences across means. Asterisk, P < 0.05; n.s P > 0.05. Longer arrows indicate cells with higher Pvalb expression. (F) Experiment 1, Pvalb, Syt2, and Nefm. (G) Experiment 2, Pvalb, Syt2, and Gabra4 (control). (H) The Neurofilament IC is observed in flash-frozen nuclei from frontal cortex. The Neurofilament (IC 25) cell-loading signal distribution across the Sst+/Pvalb+ interneuron subcluster. Left, cell-loadings displayed on subcluster tSNE plot. Right, gene-loading plots. The top 20 loading genes are shown at right.

This broad transcriptional pattern – present in almost all of the brain regions and many neuronal types – involved a large constellation of genes that underlie axonal and pre-synaptic function. We call this signal the “Neurofilament IC,” because three of most strongly contributing genes encode the neurofilament subunits of the axonal cytoskeleton (*Nefl*, *Nefm*, and *Nefh*) (Yuan et al., 2012). Other strongly co-regulated genes in this module included included *Syt2*, *Vamp1*, and *Cplx1* – which have roles in vesicle exocytosis – and *Pvalb* and *Caln1*, which bind presynaptic Ca^2+^ (**Figure S6A**). Thus, the genes nominated by this transcriptional pattern are functionally related in the maintenance of axonal function and the support or tuning of neurotransmitter release.

Neurofilament ICs were observed in all sampled brain regions and in multiple neuronal classes, including GABAergic, glutamatergic, and neuromodulatory neurons. The expression levels of genes with strongest Neurofilament IC contributions tended to covary strongly both within and across neuronal types. Among interneurons, the Neurofilament IC cell loading was most prominent in *Pvalb^+^* interneurons, which tend to have higher firing rates than other interneurons (Hu et al., 2014). For example, in frontal cortex *Sst*^+^/*Pvalb*^+^ [interneuron cluster 2](http://dropviz.org/?_state_id_=f5d0736c691e1d47), this constellation of genes exhibited continuously varying expression across *Pvalb*^+^ cells, suggesting that it contributes to variation among cells of the same subtype (**Figure 6B-D**). Among *Pvalb*^+^ interneurons, expression levels of the genes most strongly contributing to the Neurofilament IC – including [*Nefh*, *Kcnc3*, *Syt2*, and *Nefm*](http://dropviz.org/?_state_id_=0065fae0b4dff4d1) – were continuously distributed and strongly correlated with one another; by contrast, expression levels of these genes were lower and less correlated among *Sst*^+^ interneurons (**Figure 6D**). In the hippocampus, [*Pvalb*^+^ interneurons](http://dropviz.org/?_state_id_=a2f61684b3c145e4) exhibited high cell loading for the Neurofilament IC, as did the [*Pvalb*^+^ “prototypical”](http://dropviz.org/?_state_id_=940c871567f599ec) neurons of the globus pallidus externus (Mallet et al., 2012; Saunders et al., 2016)(**Figure S6A**). We also identified this axis of transcriptional variation among glutamatergic populations, for example the subplate neurons in the frontal cortex (**Figure S6A**).

To determine whether these co-expression relationships are present *in vivo*, we performed three-channel transcript counting using single-molecule fluorescent *in situ* hybridization (smFISH). The Drop-seq data predicted correlated expression of *Pvalb*, *Syt2*, and *Nefm* – but not *Gabra4* (selected as a control gene) – among *Pvalb*^+^ neurons in the frontal cortex (**Figure 6E**). This prediction was strongly confirmed by smFISH: among *Pvalb*^+^ interneurons, *Syt2* transcript densities correlated with transcript densities for *Pvalb* and *Nefm* (**Figure 6F**) but not the control gene *Gabra4* (**Figure 6G**). We did not observe any obvious spatial organization of *Syt2/Nefm* expression among *Pvalb*^+^ cells. To confirm that the Neurofilament signal did not arise from a cell-preparation artifact (such as transcriptional responses to axotomization during cell dissociation), we performed Drop-seq analysis of 28,194 single nuclei isolated from flash-frozen mouse frontal cortex; Neurofilament IC cell loading was still strongly visible among the nuclei of the *Pvalb*^+^ interneurons (**Figure 6H**).

We conclude that many neuronal types share a coordinated transcriptional program involving genes whose functions are to facilitate maintenance, expansion, or subcellular transport to the axon and presynaptic terminal. Neuronal types characterized by extensive axonal arbors, long-distance axonal projections, and/or faster firing rates tended to utilize this transcriptional program more than other neurons. At the same time, the magnitude of expression of this program varied among neurons of the same subtype, suggesting that this program contributes to both intra- and inter-type diversity.

### Gene-gene Co-expression Relationships Inferred from Hundreds of Cell types and States

Cells have diverse gene expression patterns – yet functional imperatives and regulatory architecture may greatly constrain genes’ patterns of co-expression. Gene expression data for the 565 cell populations (identified by the above subclustering analysis) make it possible to analyze genes’ co-expression relationships across many cell types, cell states and other functional contexts. We performed such analyses at the level of the 565 transcriptionally distinct cell populations (rather than 690,207 individual cells), a level of organization at which the data sample functional diversity yet are less influenced by single-cell noise and statistical sampling. (The 565 subclusters contained on average 565 cells, with expression patterns defined by an average of 1.9 million UMIs.) (**Figure S7A**).

To assess whether gene-gene correlations across the 565 cell populations could capture known functional relationships, we first focused on genes encoding subunits of the nicotinic acetylcholine receptors ([nAChRs](http://dropviz.org/?_state_id_=83f19ce3b2b27665), n=16 genes). nAChRs are ligand-gated, pentameric ion channels which we chose as a test system because the channels observed *in vivo* consist of eclectic but well-described subunit combinations that vary by brain region and neuron type (Gotti et al., 2006). We hypothesized that the observed combinations are at least partially achieved through cell-type-specific patterns of RNA expression (in addition to selective protein association).

Across the 565 cell populations, expression levels of nAChR genes exhibited two prominent correlation blocks, each containing genes that encode subunits of known heteromeric α/β channels (Zoli et al., 2015)(**Figure 7A**). For example, the expression of [*Chrna3* and *Chrnb4*](http://dropviz.org/?_state_id_=0a7e0dc2e7dd668b) (known to form a functional heteromeric receptor) were positively correlated across a large range of average transcript abundance (from 0.01 to 100 transcripts per 100K total transcripts) (**Figure 7B**). The expression of other pairs of genes that encode subunits of known heteromeric receptors ([*Chrna6/Chrnb3*](http://dropviz.org/?_state_id_=b44ad256d27d373d) and [*Chrna4/Chrnb2*](http://dropviz.org/?_state_id_=506c258f102d6205)) were also well-correlated with each other, whereas *Chrnb1* and *Chrnb2* were if anything negatively correlated, consistent with a lack of β1/β2 channels described in the brain (**Figure 7B**). These selective correlations match prominent subunit combinations inferred at the protein level *in vivo*, suggesting that nAChR composition is achieved at least in part by cell-type-specific patterns of RNA expression.

**Figure 7.**
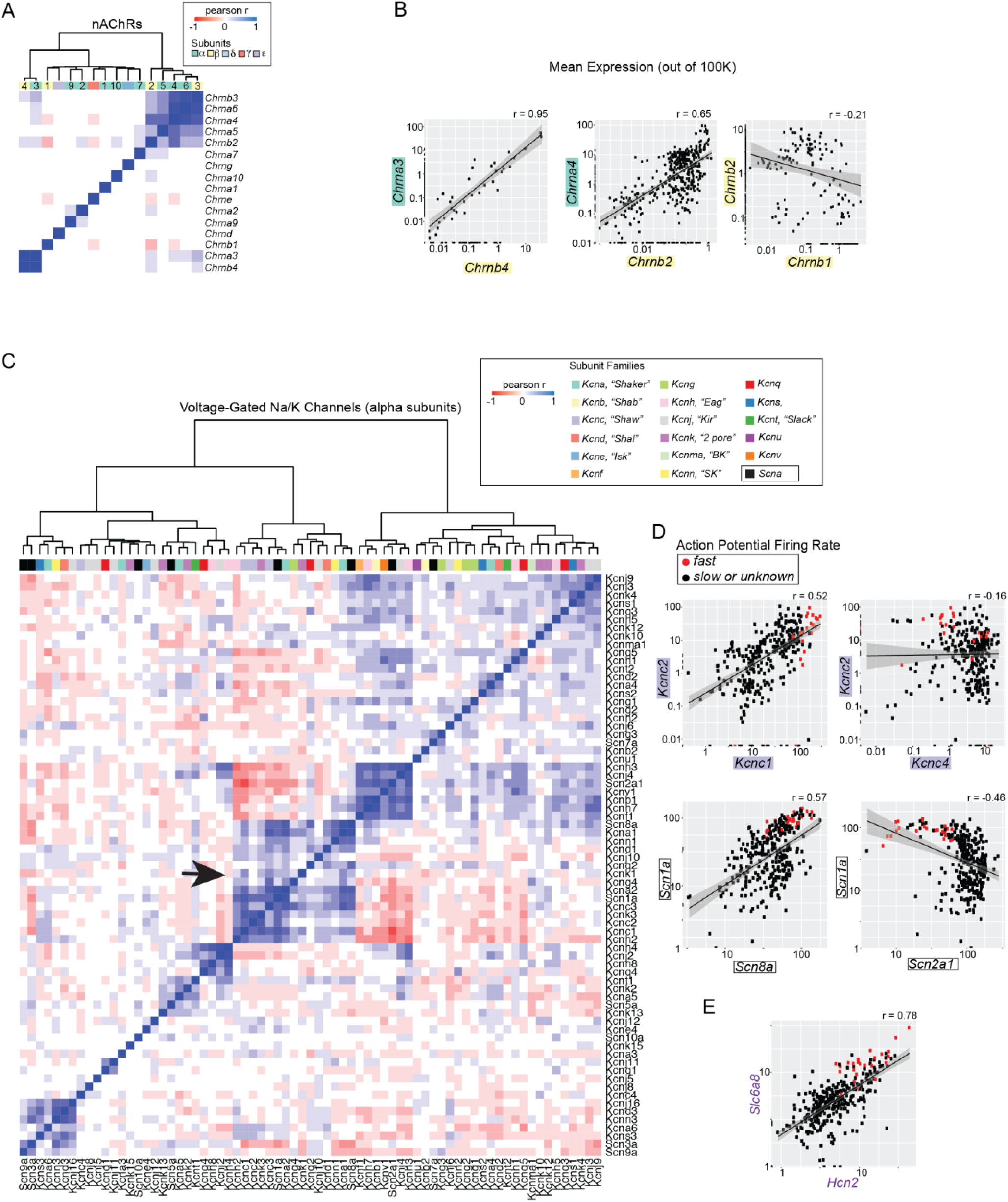
Inferring ion channel gene-gene co-expression relationships by comparing hundreds of brain cell types and states. (A-B) Nicotinic acetylcholine receptor (nAChR) subunit composition can be inferred from gene expression correlations found in 565 transcriptionally distinct atlas cell populations. Pearson correlations calculated from linear expression data. (A) Hierarchical clustering of pairwise correlations of n=16 nAChR subunit genes. Subunit families are color-coded. (B) Scatterplot comparisons of subunit expression levels. Expression levels are scaled to 100K UMIs and shown on a log10 scale. α and β subunit genes are color-coded. Note, Chrna3/Chrnb4/Chrna5 are adjacently located on chromosome 9; Chrna4/ Chrnb2 genes are on chromosome 2 and 3, respectively. (C-E) Correlation structure amongst voltage-gated (VG) Na and K channels measured from n=323 neuronal populations. (C) Hierarchical clustering of pairwise correlations calculated from expression data. The n=17 VGK channel alpha subunit families are color-coded and labeled with a common name if used. The n=1 VGNA channel alpha subunit family is shown in black. The correlation block encompassing channels known to control firing rate is shown with an arrow. (D-E) Scatterplot comparisons of subunit expression levels. Neuronal populations known to exhibit fast firing rates are shown in red (Figure S7D). Gene families are color-coded. Expression levels are scaled to 100K UMIs and shown on a log10 scale. Slc6a8 and Hcn2 were amongst the genes most frequently correlated with the alpha subunit gene set that putatively encodes firing rate (Figure S7D).

Some nAChR subunits form homomeric functional channels; the absence of obligate partners for such subunits could in principle cause them to lack positive pairwise correlations with other subunit genes. We observed several α-subunit genes with this property, including those known (*Chrna7*, *Chrna9*, and *Chrna10*) and others not yet known (*Chrna1* and *Chrna2*) to form homomeric channels in the brain (Gotti et al., 2006). In summary, gene expression correlations across cell types recapitulate known nAChR receptor subunit combinations (Gotti et al., 2006) and do so more accurately than correlations based on the bulk expression profiles for the nine regions (**Figure S7B**).

We then sought to explore the expression-correlation structure for other vital gene families whose co-function relationships are less well understood. An elegant body of experimental and theoretical work suggests that individual neurons can attain their type-specific electrophysiological properties by expressing unique combinations of voltage-gated ion channels (Marder and Goaillard, 2006). It is not known, however, if neurons can utilize any combination of ion channel genes, or face constraints (functional or regulatory) that limit which channel subunits are expressed in the same cell types. We therefore evaluated the correlations among the expression patterns of 71 voltage-gated potassium (VGK) and sodium (VGNA) channel genes across 323 neuronal populations, focusing on the genes that encode the alpha (pore-forming) subunits. This analysis identified prominent blocks of positive and negative correlation, with each such block including combinations of VGK and VGNA genes (**Figure 7D**).

To evaluate whether these gene expression– correlation blocks relate to cells’ electrophysiological properties, we identified fast-firing cell types among the 323 neuron populations (**Figure S7D**). Among two prominent sets of co-expressed genes (which were negatively correlated to each other), the larger set contained genes that enable fast and persistent action potential firing rates, including the *Kcnc1*-*3* (Kv3.1), *Kcna1* (Kv1.1) and *Scn8a* (Nav1.6) channels (Chen et al., 2008; Goldberg et al., 2008; Rudy and McBain, 2001) (**Figure 7D**). Fast-firing cell types expressed high levels of the VGK channel genes [*Kcnc2* and *Kcnc1*](http://dropviz.org/?_state_id_=808251cb3173eb85) and the VGNA channel genes [*Scn1a* and *Scn8a*](http://dropviz.org/?_state_id_=a363c3809c46e768) (the only two VGNA channels in the block). Fast-firing cells expressed variable levels of *Kcnc4* (the fourth Kv3 family member) and consistently low levels of *Scn2a1*, whose expression was inversely correlated with *Kcnc1*-*3* and *Scn1a* (**Figure 7D**). These relationships may nominate functional hypotheses about channels whose contributions to cells’ physiological properties are only partly understood. For example, *[Kcnc4](http://dropviz.org/?_state_id_=b09b120404939391)* could be a Kv3 family member that helps tailor membrane properties orthogonal to firing frequency, and *[Scn2a1](http://dropviz.org/?_state_id_=a0a1ae1d34dce3fc)* might undermine fast-firing. Interestingly, many of the genes whose expression was correlated with fast-firing properties are transcriptionally activated during postnatal maturation of fast-spiking interneurons (Okaty et al., 2009).

Finally, we asked what other genes were consistently co-expressed with the alpha subunit channel genes that themselves exhibited high expression in fast-firing neurons (**Figure S7D**). This collection contained expected genes (like those found in the Neurofilament signal, **Figure 6**), plausible ion channels (such as *Hcn2*), and unexpected genes, such as the transcription factor *[Foxj1](http://dropviz.org/?_state_id_=4eaa20623b99a24d)* and the creatine transporter *[Slc6a8](http://dropviz.org/?_state_id_=4730ae94f4ccc7c6).* Indeed, expression levels of *Slc6a8* and *Hcn2* were strongly correlated and high in fast-firing cell types (**Figure 7F**). Humans with dominant-negative mutations in *Slc6a8* often present with epilepsy, intellectual disability, and motor symptoms (van de Kamp et al., 2013), but the cellular mechanisms remain largely unknown. Our data suggest that *Slc6a8* plays an enhanced role in fast-spiking cells – which would include *Pvalb*^+^ basket cells, the resident fast-firing inhibitory neurons in cortical areas known to seed seizures. Indeed, male mice with mutations in (x-linked) *Slc6a8* have fewer GABAergic synapses (Baroncelli et al., 2017). These data highlight how analyses of gene co-expression across a large number of cell types can nominate new hypotheses about genes, brain circuitry, and disease.

### Cell-type Specialization Between Cortical Poles

The cerebral cortex processes motor, sensory, and associative information and is expanded in primates, especially humans (Buckner and Krienen, 2013). While extrinsic inputs are specific to different cortical areas, little is known about what intrinsic molecular specializations could contribute to region-specific cortical function. We first determined how accurately our cortex datasets represents cellular populations *in vivo* and then systematically identified transcriptional specializations within each non-neuronal cell class and across the two major neuronal types (glutamatergic and GABAergic neurons).

The process of single-cell dissociation followed by the capture, amplification, and sequencing of single-cell transcriptomes may distort the abundance of brain cell types. To test for distortions in our tissue preparation or analysis, we compared the abundances of molecularly-defined cell populations in our frontal cortex dataset to intact tissue using immunohistochemistry and *in situ* hybridization (**Figure S8D,E** and **Methods**). We found that neurons were over-represented relative to non-neurons in the Drop-seq dataset ([Drop-seq](http://dropviz.org/?_state_id_=ecc38745e333eaa8): 0.76 ± 02 mean ± sem; tissue: 0.51 ± 03). We believe this effect is partially explained by cell-inclusion thresholds used for analysis, in which some real single-cell libraries are not included due to their small size; neurons, which had over three-fold higher transcript counts than non-neurons, are relatively protected from this effect (neurons: 5,039 ± 15 mean ± sem; non-neurons: 1,696 ± 9, **Figure S8F** and **Methods**). Compared to GABAergic interneurons, glutamatergic neurons exhibited two-fold greater abundance than expected from tissue (ISH: 5.1:1; [Drop-seq](http://dropviz.org/?_state_id_=6b28d5d18d69e4df): 11:1)(Sahara et al., 2012), an effect that again could be driven by larger libraries (glutamatergic: 5,298.916 ± 16, mean ± sem; GABAergic: 2,626 ± 21) (**Figure S8H**). Comparing [three major subtypes](http://dropviz.org/?_state_id_=7b7301b516d6253b) of GABAergic interneurons, we observed that *Vip^+^* cells were overrepresented and *Sst*^+^ and *Pvalb*^+^ cells were underrepresented in the data (*Vip*^+^ ISH: 16%; Dropseq: 35%; *Pvalb*^+^ ISH: 31%; Dropseq: 25%; *Sst*^+^ ISH: 28%; Dropseq, 22%). Since greater *Vip*^+^ interneuron abundance cannot be explained by higher transcript counts (*Pvalb*^+^: 2,996 ± 61; *Sst*^+^: 2,758 ± 53; *Vip*^+^: 2,236 ± 32), these data may suggest that *Pvalb/Sst* interneurons were preferentially depleted during the preparation. This distortion could also contribute to the increased fraction of glutamatergic/GABAergic neurons that we observe. We conclude that our data exhibit modest, quantitative skews in cellular representation that are primarily driven by differences in transcript abundance and cell viability.

To identify molecular specializations across cortical regions, we performed semi-supervised ICA on cells from the frontal and posterior cortex together, separately analyzing cells of the various cell classes: excitatory neurons, inhibitory neurons, astrocytes, oligodendrocytes/polydendrocytes, microglia/macrophages, and fibroblast-like and endothelial cells. To identify transcriptional signals that differed between regions, we first calculated a “region skew score” for each of the biological ICs (based on the tendency of high-scoring cells to come from one region or the other, relative to low-scoring cells) (**Figure 8A, Methods**). Glutamatergic excitatory neurons generated more such regionally specialized ICs than GABAergic interneurons and non-neuronal cell classes did. Consistent with this, we identified strong regional differences in cellular abundance (a normalized ratio of greater than 3:1, P < 0.05, Barnard’s Test, Bonferroni corrected) for seven subclusters of excitatory neurons and none of the other cell classes (**Figure 8B,C and Figure S9A**). To visualize the spatial distributions of the excitatory-neuron populations corresponding to these subclusters, we examined the ISH patterns of markers for each of these populations; the ISH patterns consistently confirmed asymmetric expression across the cortical mantle (**Figure 8C and Figure S9B**). To determine whether molecular differences, in addition to differences in cell-type representation, contributed to frontal versus posterior specialization, we performed differential expression analysis comparing cells (within the same subcluster) from the two cortical regions. These analyses revealed far more differentially expressed genes in comparisons of excitatory populations than in comparisons of GABAergic interneurons or non-neuronal cell classes (**Figure 8D**). These data suggest that molecular specializations across cortical areas are largely driven by differences involving excitatory projection neurons. This result is consistent with independent single-cell analyses in mouse (Tasic et al., 2017) and with theories of cortical specialization in humans and other primates (Krienen et al., 2016).

**Figure 8.**
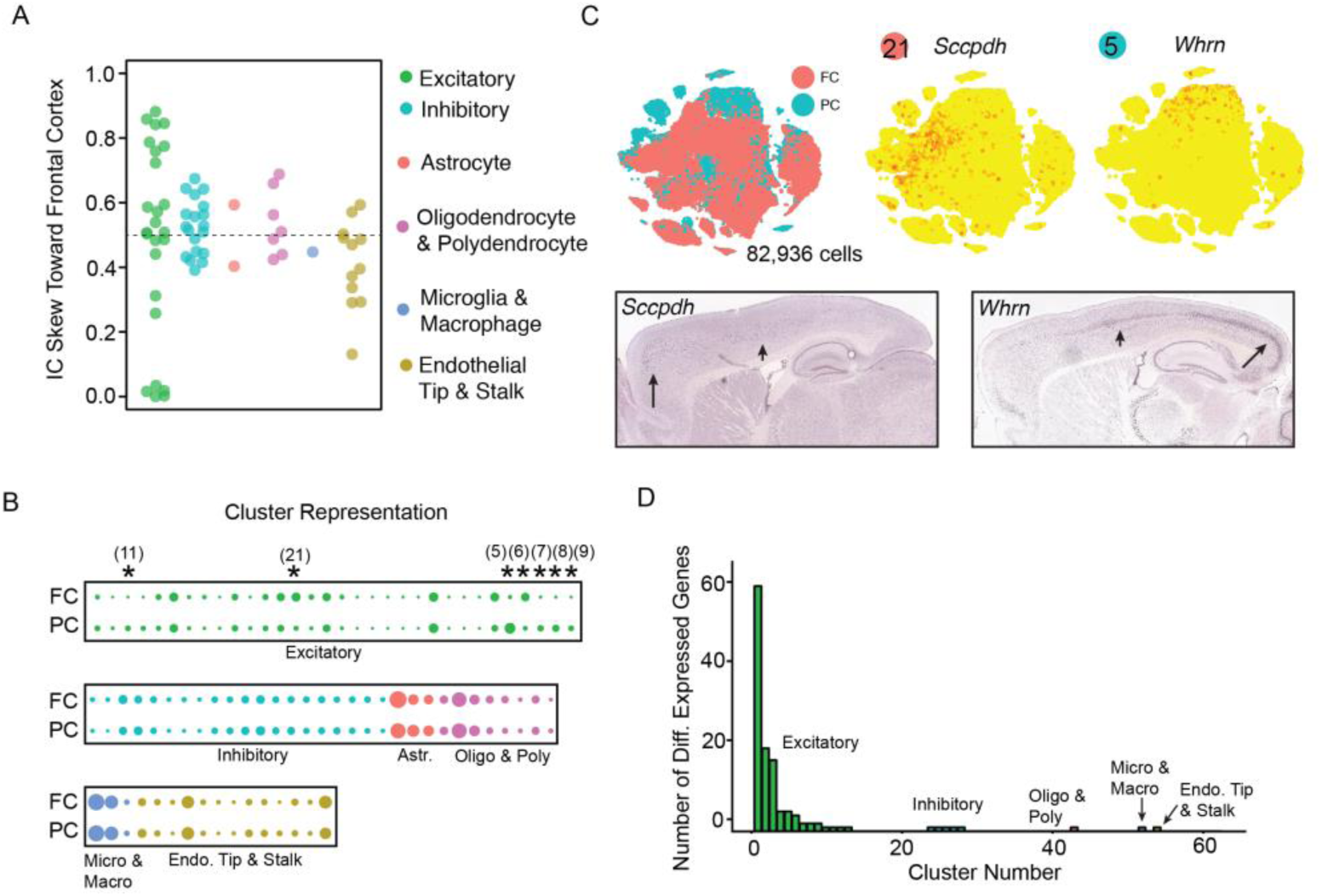
Cortical regions remain transcriptionally specialized via excitatory glutamatergic neurons. A. Beeswarm plot showing the relative contribution of frontal (FC) vs posterior (PC) cells to top-loading cells for each of the biological ICs in six separate cell-class analyses (glutamatergic excitatory neurons, n=82,936 cells; GABAergic inhibitory neuron, n=7,783; astrocyte, n=7,782, oligodendrocytes/polydendrocytes, n=3,505; microglia/macrophage, n=1,027; and endothelial/FL cells, n=3,578). For each analysis, equivalent cell classes from the frontal and posterior datasets were jointly reanalyzed. The normalized proportion of cells in FC and PC is plotted. The IC Skew score is 1 if only FC cells contribute and 0 if only PC cells contribute. Equal contribution is 0.5 (dotted line).
B. Plot comparing the relative number of FC vs PC cells within each subcluster following clustering of the biological ICs for each cell class analysis. Circle size denotes the fractional representation. Subclusters with greater than 3:1 compositional skew are indicated with an asterisk and their label is shown (P < 0.05, Barnard’s test, Methods). Subclusters plots for each cell class are shown in Figure S9.
C. Subcluster analysis of glutamatergic excitatory neurons from FC and PC illustrating that excitatory neuron populations are transcriptionally specialized by cortical region. Top left, tSNE plot of excitatory neurons color-coded by region. Top right, expression tSNE plots for Sccpdh and Whrn, genes enriched in subclusters 21 and 5 that are disproportionately composed of FC or PC cells respectively. Bottom, ISH stain (Allen Institute Mouse Brain Atlas) for Sccpdh and Whrn, which exhibit skewed expression in the FC or PC, respectively. High expression, long arrow; Medium expression, short arrow.
D. Barplot summarizing the number of genes differentially expressed across FC and PC cells within each subcluster across all cell class experiments (> 2-fold change, P < .05, Bonferroni corrected).

### Resolving Neuron Types within the Basal Ganglia

Neuronal types and subtypes in many brain regions are incompletely identified or characterized, limiting their study in neural circuits, behavior, and disease. Our data provide an opportunity to more systematically identify neuronal types in many regions. From the 9 regions, we identified [323 neuronal subclusters](http://dropviz.org/?_state_id_=bc20d0f2e4f7ddf0) which arose from 368 biological ICs (**Figure S4**).

Here, we illustrate the identification of neuronal types by focusing on the basal ganglia, a collection of richly interconnected subcortical areas. While neuron types in the striatum have received most attention, decades of *in vivo* and *in vitro* electrophysiological recordings in other basal ganglia nuclei have documented extensive neuronal diversity – yet the anatomical complexity of these regions has made identifying neuron types challenging and laborious (Bolam et al., 2000; DeLong et al., 1984). We turned to the globus pallidus externus (GPe) and substantia nigra reticulata (SNr), two highly disease-relevant (Albin et al., 1989) regions in which neuron types are not yet well classified (**Figure 1 & Figure S1**).

Like many functionally specialized nuclei in the rodent brain, the GPe and SNr are anatomically small structures, with 10^3^–10^5^ total neurons (Oorschot, 1996), as compared to the 10^6^–10^7^ found in larger regions like the striatum, cortex, and hippocampus (Abusaad et al., 1999; Schüz and Palm, 1989). This poses a challenge for Drop-seq, which samples only about 5% of the total cells in a cell suspension (Macosko et al., 2015). We addressed this issue by both optimizing dissociation (to maximize cell yield, **Methods**) and sampling larger volumes of tissue that encompassed multiple adjacent structures. (We note, however, that as the number of included regions goes up, so too does the chance for biases in neuronal representations (because of differences in library size and sensitivity to the digest), as well as the complexity and ambiguity of the post-hoc anatomical analysis.) The GPe and SNr were included in the GP/NB and SN/VTA analyses, respectively (**Figure S1**).

To identify GPe neuron populations, we leveraged the Allen Institute’s digital ISH atlas to spatially map expression of markers of global clusters and subclusters to the GPe or surrounding regions from the GP/NB dataset (**Figure 9, Figure S1**). GPe neurons were present in [cluster 2](http://dropviz.org/?_state_id_=9976d7061548b632) (n=11,103 cells), one of three neuron clusters we identified. [Cluster 1](http://dropviz.org/?_state_id_=b071d3deb936860c) corresponded to cholinergic neurons from the NB and GPe (n=437 cells) and [cluster 3](http://dropviz.org/?_state_id_=06980750fda8f838) to neurons of the adjacent striatum and basolateral amygdala (n=9,847 cells; **Figure 9A**).

**Figure 9.**
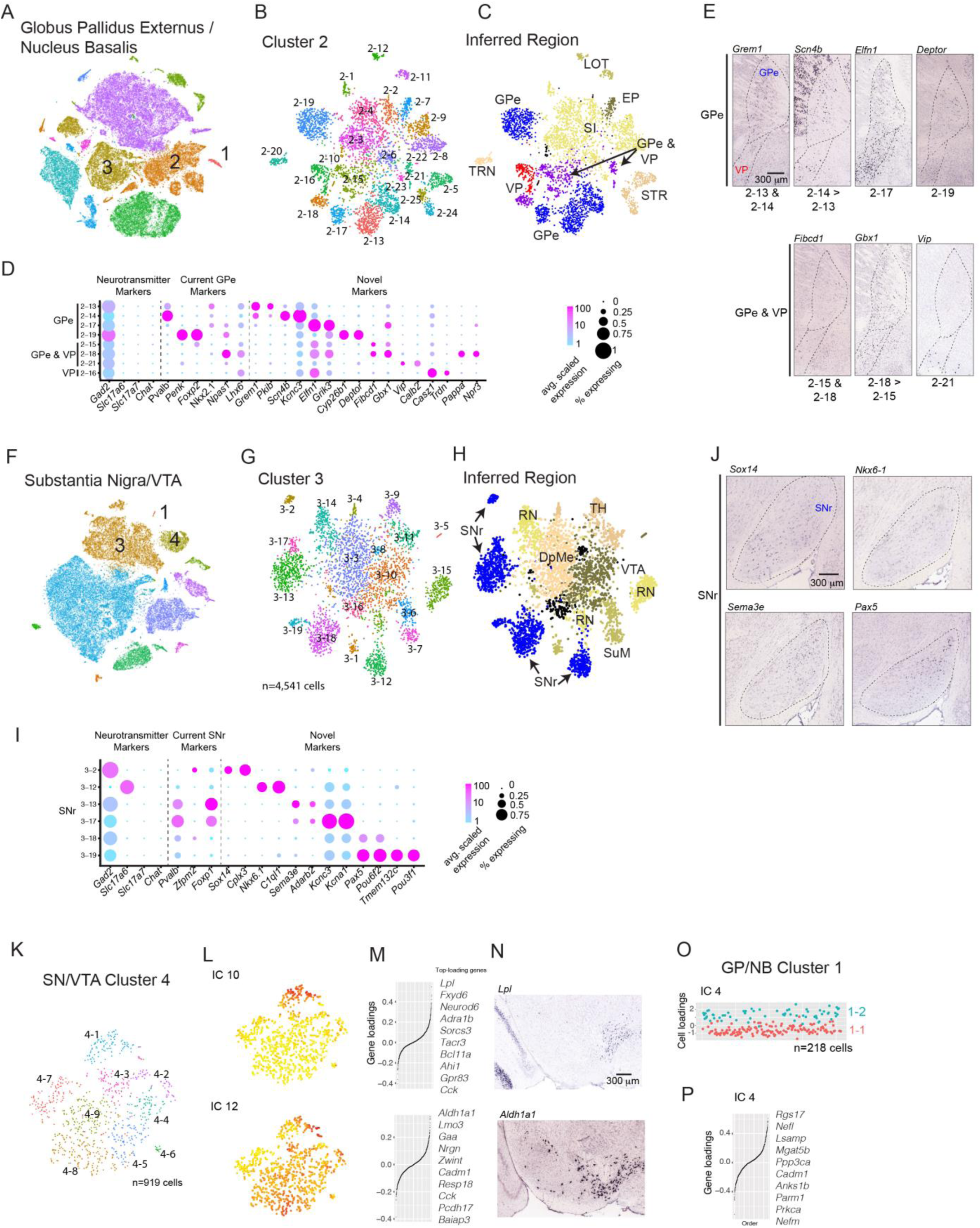
Transcription-based identification of known and novel neuron type distinctions within the basal ganglia. (A-E) Globus pallidus externus (GPe). (F-J) Substantia nigra reticulata (SNr). (K-O) Dopaminergic vs acetylcholinergic neuromodulatory neuron populations. (A) Color-coded global clusters (n=11) for globus pallidus externus/nucleus basalis dataset displayed on a tSNE plot. Clusters 1, 2, and 3 are neuronal. (B) Subcluster structure within cluster 2. (C) Subclusters color-coded by candidate anatomical regions, inferred by ISH expression patterns of selective marker genes (Figure S10). Populations are intrinsic to the GPe, ventral pallidum (VP), substantia innomatita (SI), adjacent striatum (STR), lateral olfactory tract (LOT), rostral entopeduncular nucleus (EP) and the thalamic reticular nucleus (TRN) consistent with dissections (Figure S1). (D) Dot plot illustrating the expression patterns of neurotransmitter marker genes, neuron type markers from the literature, and novel markers identified in this analysis. Dot diameter represents the fraction of cells within a subcluster where a transcript was counted. Colors represent average single-cell scaled expression value (out of 100K UMIs, log10). (E) ISH experiments with subsets of novel marker genes illustrating the expression within the anatomical regions of the GPe and/or VP (sagittal sections). Region borders are approximated by a dotted line (Allen Institute Mouse Atlas). (F) Color-coded global clusters (n=14) for substantia nigra/VTA dataset displayed on a tSNE plot. Clusters 1, 3, and 4 are neuronal. (G) Subcluster structure within cluster 3. (H) Subclusters color-coded by candidate anatomical regions, inferred by ISH expression patterns of selective marker genes (Figure S10). Populations are intrinsic to the SNr, ventral tagmental area (VTA), red nucleus (RN), supramammillary nucleus (SuM), thalamus (TH), and deep mesencephalic nucleus (DpMe). (I) Dot plot as in (D). Marker genes for neurotransmitters, current SNr markers, and novel markers identified in this study. (J) ISH experiments with subsets of novel marker genes illustrating the expression within the anatomical regions of SNr (sagittal sections). Region borders are approximated by a dotted line (Allen Institute Mouse Atlas). (K) Subcluster structure within Th+/Ddc+ dopaminergic cluster 3 from the SN/VTA dataset. (L-M) Example cluster 3 ICs that encode spatial signals within the SNc/VTA. (L) IC cell loadings displayed on tSNE plot. (K) IC gene-loadings. Top ten genes shown at right. (L) ISH experiments (sagittal sections) for Lpl (IC 10, top) and Aldh1a1 (IC 12, bottom). IC 10 identifies the dorsal VTA, while IC 12 identifies the ventral VTA and SNc (Allen Institute Mouse Atlas). (O-P) Minimal heterogeneity identified within Chat+/Slc5a7+ cholinergic cluster 1 from the GP/NB dataset. (O) Plot of IC 4 cell-loadings. Based on IC 4, cells are assigned as subcluster 1-1 or 1-2. (P) IC 4 gene-loading plot. Top ten loading genes at right suggest a Neurofilament-type signal.

To determine which of the [25 subclusters](http://dropviz.org/?_state_id_=d395711903d66ab2) within cluster 2 were intrinsic to the GPe or the adjacent ventral pallidum (VP), we identified markers from our data and evaluated their spatial expression using ISH. We found [8 candidate subclusters](http://dropviz.org/?_state_id_=8eee53eea4e26fbf) intrinsic to the GPe and/or VP, of which 4 were [exclusive to the GPe](http://dropviz.org/?_state_id_=7b1ee175de5b7f39) (**Figure 9C-E**). The other 17 subclusters mapped to surrounding regions, including the thalamic reticular nucleus (TRN), the substantia innominata (SI), and the lateral olfactory tract (LOT) (**Figure S10**).

Modern approaches to classify GPe neurons rely on iterative mapping of anatomical, electrophysiological, and behavioral response properties onto single molecular markers (e.g., (Mallet et al., 2012; Saunders et al., 2016)); such data provide essential insights into neuronal diversity, but have trouble conclusively identifying distinct populations. To associate subclusters with putative GPe neuron types, we compared the expression of published GPe markers to pairs of markers that we identified for each subcluster (**Figure 9D**). Of [markers described previously](http://dropviz.org/?_state_id_=abe918da8b5fc1d1), only *Pvalb* and *Penk/Foxp2* were selectively expressed in particular GPe subclusters (Kita, 1994; Voorn et al., 1999). This correspondence suggests subcluster 2-14 represents the fast-spiking “prototypical” population, while 2-19 represents the slow-firing “arkypallidal” population (Abdi et al., 2015; Mallet et al., 2012). Interestingly, 2-13 is transcriptionally similar to 2-14, sharing markers such as *[Grem1](http://dropviz.org/?_state_id_=ffed00c8e3922a29)*, but is [distinguished](http://dropviz.org/?_state_id_=c6de3551e10b1025) by its stronger expression of genes like [*Scn4b* and *Kcnc3*](http://dropviz.org/?_state_id_=8d49a3fcccc8dc91) and those from the “Neurofilament program” (described above) (**Figure S6A**). Further experiments will be necessary to determine whether subclusters 2-14 and 2-13 represent anatomically distinct subpopulations or transient states. The fourth GPe subcluster (2-17) is enriched for markers *[Elfn1/Grik3](http://dropviz.org/?_state_id_=0a8d584a242ae544)*, but to our knowledge, has not been characterized. Other published markers (e.g., *Nkx2.1*, *Npas1*, and *Lhx6*) show mild, reciprocal enrichments across multiple subclusters, illustrating how markers can fail to capture the true population structure of neurons (Abdi et al., 2015; Hernandez et al., 2015; Mastro et al., 2014).

To our surprise, several subclusters (2-15, 2-21, and 2-18) expressed markers found in both the VP and bordering GPe (**Figure 9D,E**). This border-spanning ISH pattern suggests that some neuron types may be found in regions with differing behavioral functions and connectivity (Gittis et al., 2014; Kita, 2007; Smith et al., 2009). Conceptually, this may be analogous to distinct regions of the cerebrum harboring similar interneurons (Harwell et al., 2015). These shared subclusters may explain a neuron type that is synaptically incorporated into the GPe, but exhibits VP-like axonal projections (Chen et al., 2015b; Saunders et al., 2015).

To identify the neuron populations intrinsic to the SNr, we followed a similar procedure using the SN/VTA dataset (**Figure 9F, Figure S1**). Of the [4 global neuron clusters](http://dropviz.org/?_state_id_=00966e21874f5a02) identified, one (cluster 3, n=10,049 cells) contained neurons of the SNr as well as the surrounding regions (**Figure 9H**). The other three clusters (clusters 1, 2, and 4, respectively containing 73, 297, and 1,841 cells) contained hippocampal, thalamic, and dopaminergic neurons, respectively.

Of the [19 subclusters](http://dropviz.org/?_state_id_=2a6eda739dbbb750) in cluster 3, we mapped [6 to the SNr](http://dropviz.org/?_state_id_=87a9f6c46276c2a6) (**Figure 9G,H** and **Figure S10B**). Two sets of subclusters shared highly [selective markers](http://dropviz.org/?_state_id_=187a145dbca39ac4) (3-18/3-19: *Sema3a*, *Adarb2*; 3-17/3-13: *Pax5*, *Pou6f2*), suggesting they are closely related. Each pair is distinguished by genes whose expression might imply a state or subtype distinction, like ion channels (*Kcnc3*, *Kcna1*), transmembrane proteins (*Tmem132c*), and transcription factors (*Pou3f1*) (**Figure 9I**). Of the remaining 2 SNr subclusters, 3-12 expressed *[Slc17a6^+^](http://dropviz.org/?_state_id_=ec1ce80a354c2e71).* presumably corresponding to the glutamatergic projection from the SNr to the thalamus (Antal et al., 2014). Subcluster 3-2 was *Gad2*^+^/*Pvalb*^-^, [expressed](http://dropviz.org/?_state_id_=d828ca57b0359f86) the developmental marker *Zfpm2* (Lahti et al., 2016), and likely represents a third GABAergic type marked by *Sox14* and *Cplx3* expression (**Figure 9I**). To our knowledge, this is the first comprehensive molecular description of adult SNr neurons.

The GPe and SNr each abut small populations of neuromodulatory neurons that have profound effects on circuit physiology and behavior through the widespread release of acetylcholine (ACh) and dopamine (DA), respectively. Molecular specializations are known to exist in the DA system that are, at least in part, associated with the anatomical location of DA neuron subtypes (Lammel et al., 2008; Poulin et al., 2014). It is unclear whether the ACh-releasing population exhibits similar heterogeneity and whether this diversity has a spatial component.

Subcluster analyses of the dopaminergic (SN/VTA, [cluster 4](http://dropviz.org/?_state_id_=fffc6e3698233906), 919 cells) and cholinergic (GP/NB, [cluster 1](http://dropviz.org/?_state_id_=95941b18f69c2198), n=218 cells) clusters revealed that the sampled dopaminergic neurons were indeed more heterogeneous than the cholinergic neurons (DA: 6 biological ICs, 9 subclusters; ACh: 2 biological ICs, 2 subclusters)(**Figure 9A,F**). As expected, aspects of DA transcriptional diversity were related to spatial positioning (Poulin et al., 2014) – for example, delineating the dorsal (IC 10) from ventral (IC 12) VTA as illustrated by ISH stains for the weighted genes [*Lpl* and *Aldh1a1*](http://dropviz.org/?_state_id_=feb490a56f77eb1c) (**Figure 9L-N**). In contrast, the major signal in cholinergic neurons (IC 4) appears to be Neurofilament-like, with no spatial component (**Figure 9O,P**). Thus, DA neurons of the SNc/VTA appear to be regionalized, whereas cholinergic neurons of the GPe/NB are not. Sampling more cholinergic neurons, especially from other areas of the basal forebrain (Zaborszky et al., 2013), could reveal additional signals. Spatial transcriptomic technologies that measure single cells could also reveal anatomical relationships missed by ISH analysis on thin slices of tissue (Chen et al., 2015a; Shah et al., 2016).

### Molecular Specializations of Striatal Principal Neurons

Spiny projection neurons (SPNs) represent about 95% of the neurons in the rodent striatum. Decades of anatomical and functional work has sought to establish how molecularly defined subtypes of SPNs contribute to circuit function, behavior, and disease (Albin et al., 1989; Kozorovitskiy et al., 2012; Kravitz et al., 2010). Two principal categories have been used to distinguish SPN subsets. The first – based on divergent axonal projections and receptors for dopamine – assigns SPNs to the “direct” (dSPN) and “indirect” (iSPN) pathways; dSPNs and iSPNs are similarly numerous. The second – based traditionally on processing limbic versus sensory/motor information – groups SPNs into two spatial compartments within the striatum, the so-called “patch” and “matrix” compartments (Gerfen, 1992; Graybiel and Ragsdale, 1978). Both dSPNs and iSPNs are present in the patch and matrix.

The direct- and indirect-pathway SPN types were readily identified by the expression of the pan-SPN marker *Ppp1r1b* along with the iSPN marker *Adora2a* or the dSPN marker *Drd1* (**Figure 10A,B**). Two large clusters were highly enriched for *Ppp1r1b/Drd1* and *Ppp1r1b/Adora2a*, suggesting they corresponded to dSPNs (Cluster 10, n=30,835 cells) and iSPNs (Cluster 11, n=25,305 cells), respectively. There were 68 [differentially expressed genes](http://dropviz.org/?_state_id_=b89ec7ffb19dd21d) between the two, including both known and previously undescribed markers (**Figure 11A** and **Table S2**).

**Figure 10.**
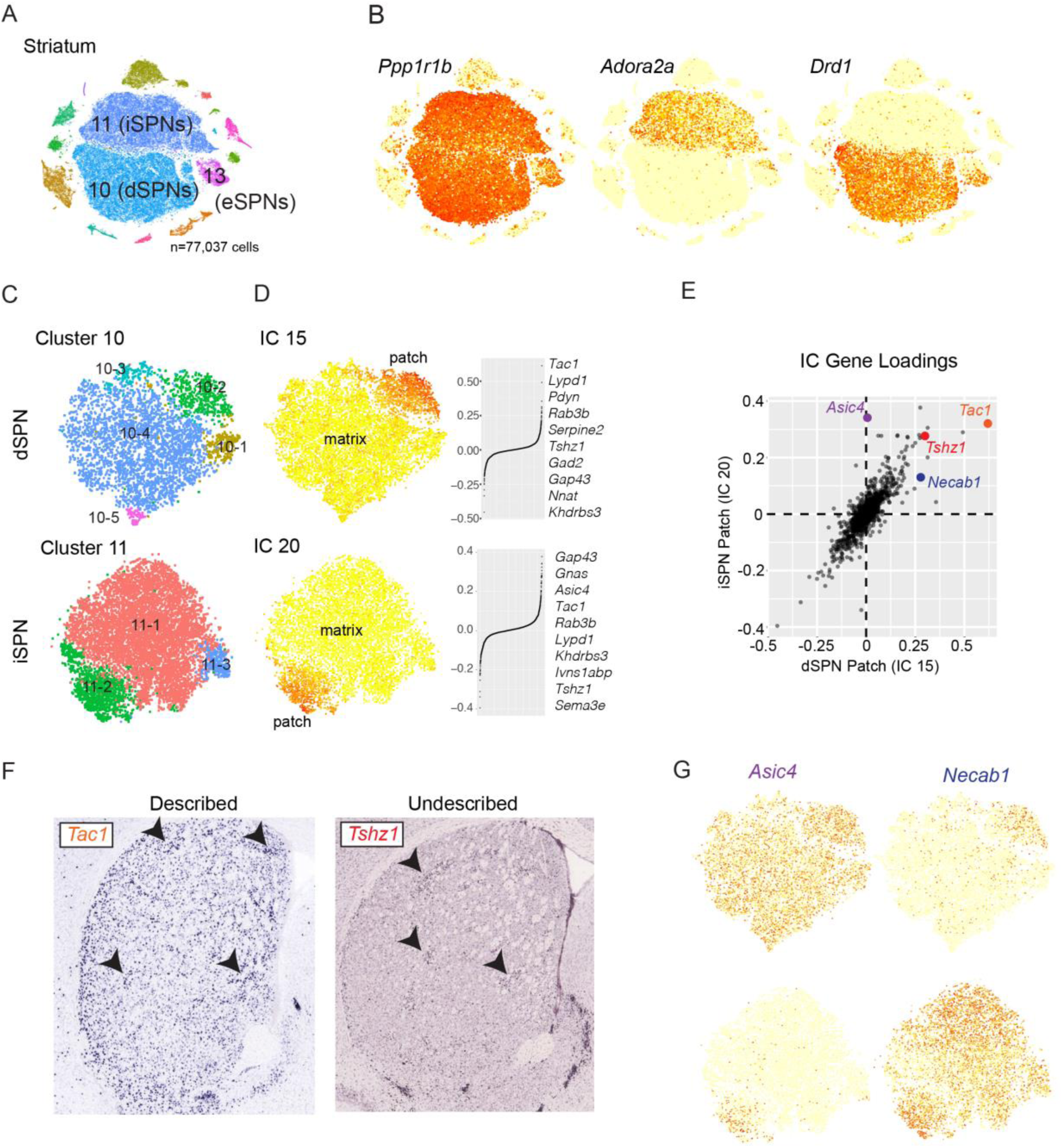
Molecular specializations within spiny projection neuron patch and matrix spatial compartments. (A) Color-coded global clusters (n=15) for striatum dataset displayed on a tSNE plot. Clusters 10, 11, and 13 are presumed SPNs, corresponding to dSPNs, iSPNs, and eSPNs (described in Figure 8). (B) Expression tSNE plot of pan-SPN marker Ppp1r1b, direct pathway SPN (dSPN) marker Drd1, and indirect pathway SPN (iSPN) marker Adora2a. Cluster 13 expresses SPN markers. Ppp1r1b+ cells within Cluster 13 are eccentric SPNs (eSPN). (C-E) Identification and comparison of patch iSPN and dSPN transcriptomes. (C) Color-coded subcluster assignments for striatum cluster 10 (dSPNs) and cluster 11 (iSPNs) displayed on tSNE plot. (D) ICs encoding patch-specific transcriptional signals in dSPNs (IC 15) and iSPNs (IC 20). Left, IC cell-loadings on tSNE plots. Right, gene-loadings with top 10 genes displayed. (E) Identification of shared and distinct dSPN and iSPN patch-enriched genes. Scatterplot comparing gene loadings between patch ICs for dSPNs (IC 15) and iSPNs (IC 20). Genes with high loading in both ICs are hypothesized to be enriched in both patch iSPNs and dSPNs. This group includes described (Tac1) and previously undescribed (Tshz1) genes enriched in patch dSPNs and iSPNs. Genes that load strongly onto either IC are candidates for patch molecular specialization by SPN pathway. For example, Asic4 is enriched in patch iSPNs, while Necab1 is enriched in patch dSPNs. (F) ISH stains for Tac1 and Tshz1 on coronal sections of striatum (Allen Mouse Brain Atlas). Arrowheads point to example patches. (G) Expression tSNE plots for Asic4 and Necab illustrating patch enrichment exclusive to iSPNs and dSPNs, respectively.

**Figure 11.**
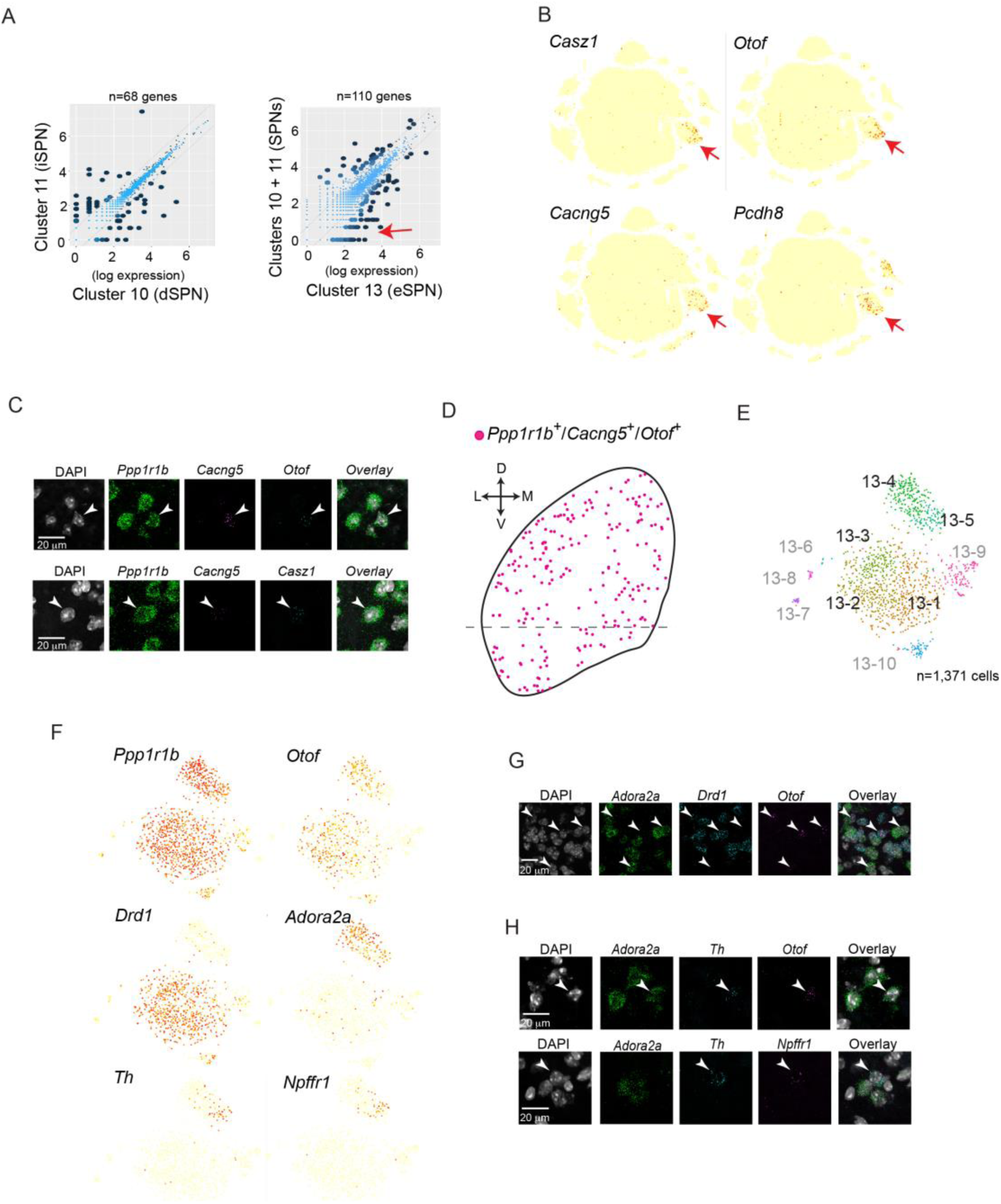
Eccentric spiny projection neurons represent a third axis of SPN diversity. (A) Genome-wide mean expression comparisons between SPN populations (log-normal scale). Left, cluster 10 vs cluster 11 (iSPN vs dSPNs). Right, cluster 13 vs clusters 10 and 11 (eSPNs vs d/iSPNs). Differentially expressed genes are shown with larger, dark dots (>2 fold difference and P<10^-100^, binomTest (Robinson et al., 2010) and total number listed above each plot. Red arrow indicates genes with selective expression in eSPNs vs d/iSPNs. (B) Expression tSNE plot of n=4 genes (*Casz1*, *Otof*, *Cacng5*, and *Pcdh8*) enriched in eSPNs vs d/iSPNs (red arrow in C). Across all global clusters, genes are highly enriched in cluster 13 (red arrows). (C-D) eSPNs are anatomically dispersed throughout the striatum. (C) Single confocal planes from smFISH experiments validating co-expression of pan-SPN (*Ppp1r1b*) and highly-selective eSPN markers (*Cacng5*, *Otof*, and *Casz1*) in dorsal striatum. Top, *Ppp1r1b*, *Cacng5*, and *Otof.* Bottom, *Ppp1r1b*, *Cacng5*, and *Casz1.* Arrowhead indicates triple-positive cells. (D) Locations of triple positive *Ppp1r1b*, *Cacng5*, and *Otof* cells on a schematic of coronal striatum. Only dorsal striatum was used in the Drop-seq analysis (approximated by the dotted line). D, dorsal; V, ventral; L, lateral; M, medial. (E) Color-code subclusters from cluster 13. Subclusters 13-1, 13-2, 13-3, 13-4, and 13-5 correspond to eSPNs (83% of cluster 13 cells, black labels). The anatomical identity of other subclusters (17% of cluster 13 cells, gray labels) is described in **Figure S12**. Non-eSPN subclusters were excluded from the differential expression analysis described in (A). (F) Expression tSNE plot of pan-SPN (*Ppp1r1b*), pan-eSPN (*Otof*), dSPN (*Drd1*), iSPN (*Adora2a*), subcluster 13-5 (*Th*, *Npffr1*) markers. (G-H) Single confocal planes from smFISH experiments validating co-expression of markers in dorsal striatum. Arrowhead indicates triple-positive cells. (G) Co-expression of eSPN marker (*Otof*) with iSPN (*Adora2a*) and dSPN (*Drd1*) markers. (H) Co-expression of eSPN subcluster 13-5 markers. Triple-positive cells observed in dorsal striatum are indicated with white arrowheads. Top, *Adora2a*, *Th*, *Otof.* Bottom, *Adora2a*, *Th*, *Npffr1.*

To identify cells from the patch and matrix compartments among dSPNs and iSPNs, we inspected transcriptional signals identified from SPN subclustering (dSPN Cluster 10: n=9 biological ICs; iSPN Cluster 11: n=10 biological ICs, **Figure S11**). Each analysis identified a single candidate “patch IC” whose most strongly contributing genes included known patch markers such as *Tac1* and *Pdyn.* Approximately 10% of iSPNs and dSPNs exhibited this patch transcriptional phenotype (dSPN: IC 15, subcluster 10-2; iSPN: IC 20 in subcluster 11-2; **Figure 10C,D**).

To appreciate the extent to which the patch/matrix distinction affects the functional specializations of dSPNs and iSPNs, we compared gene loadings between the patch-encoding dSPN IC 15 and iSPN IC 20 (**Figure 10E**). As expected, we observed both classic (*Tac1*) and undescribed (*Tshz1*) [pan-patch](http://dropviz.org/?_state_id_=578cc1677273699c) markers (**Figure 10F**). We found that the proton-gated Na channel *[Asic4](http://dropviz.org/?_state_id_=2feb1a0072cad341)* is enriched in iSPN patch cells, while Ca-binding protein *[Necab1](http://dropviz.org/?_state_id_=2feb1a0072cad341)* is enriched in dSPN patch cells (**Figure 10G**). While studies of SPNs have focused independently on the patch/matrix or pathway axes, our data suggest a more complex pattern of specialization between these systems: the transcriptional features endowed by a patch habitat are not identical for dSPNs and iSPNs, and some of these differences appear to remove pathway expression differences found in the matrix dSPNs and iSPNs. For example, among matrix SPNs, *Asic4* was selectively expressed in the dSPNs but not iSPNs; among patch SPNs, however, *Asic4* was expressed in both dSPNs and iSPNs. Thus *Asic4* helps tailor iSPNs but not dSPNs to the patch habitat.

### “Eccentric” SPNs: A novel, third axis of SPN diversity

Surprisingly, about 4% of SPNs (*Ppp1r1b*^+^) were observed in a [third, smaller cluster](http://dropviz.org/?_state_id_=8deeb6a433bc345e) that also expressed *Adora2a* and *Drd1* (cluster 13: n=2,744 cells; 4.5% of *Ppp1r1b*^+^ neurons; **Figure 10A,B**). All three biological replicates contributed to this distinct population, suggesting it was not due to a dissection artifact. These SPNs had a transcriptional phenotype distinct from other SPNs: some 110 genes differed significantly in expression between this [cluster and the canonical](http://dropviz.org/?_state_id_=8d0e82549d6039d9) dSPNs+iSPNs (fold ratio > 2 and P < 10^-100^ by binomTest (Robinson et al., 2010)), more than the number of genes that distinguish dSPNs and iSPNs from each other (n=68) (**Figure 11A, Table S3**), Several genes were selectively expressed in cluster 13 with very little expression in the rest of the striatum (e.g., *[Casz1, Otof, Cacng5, and Pcdh8](http://dropviz.org/?_state_id_=a3d75da5b738ab8b)*) (**Figure 11B**). Due to their expression of canonical SPN genes (e.g., *Ppp1r1b*) yet large transcriptional divergence from canonical SPNs, we call this population “eccentric” SPNs (eSPNs).

To determine where eSPNs are located, we performed triple smFISH labeling with *Ppp1r1b* and pairs of selective eSPN markers (**Figure 11C,D**). Candidate eSPNs (triple-positive cells) clearly localized to the striatum; like dSPNs and iSPNs, eSPNs exhibited no obvious spatial organization within the striatum and were intermixed with other SPNs. We conclude that eSPNs are an intrinsic striatal neuron population. Our Drop-seq data account for all known [striatal interneuron types](http://dropviz.org/?_state_id_=674d39074f33b7c8) at approximately the expected proportions (3.9% of total neurons)(Tepper and Bolam, 2004), suggesting by exclusion that the subclusters we identify as eSPNs do not correspond to an interneuron type but rather are SPNs.

To establish whether the eSPNs harbor additional molecular diversity, we examined the [five subclusters](http://dropviz.org/?_state_id_=39c454601e4bd7c2) in which pan-SPN (*Ppp1r1b*) and pan-eSPN (*Otof*) markers were expressed (**Figure S12A-C**). These subclusters were divided into two major groups of eSPNs (**Figure 11F. Table S4**), separated by a [gene set](http://dropviz.org/?_state_id_=71bde98c16330e6e) that included markers used to distinguish canonical iSPNs and dSPNs, such as *Drd1* and *Adora2a*, which we confirmed were expressed in eSPNs with smFISH (**Figure 11G**). eSPN expression of markers typically associated with canonical SPNs suggests eSPNs have been molecularly “camouflaged,” including in studies using mice that have employed *Drd1* and *Adora2a* driven–transgenes to label and manipulate dSPNs or iSPNs (Cui et al., 2014; Kozorovitskiy et al., 2012; Kravitz et al., 2010; Oldenburg and Sabatini, 2015). Despite sharing markers, *Adora2a*^+^ eSPNs and *Drd1*^+^ eSPNs are distinguished from their canonical SPN counterparts by expression levels of many genes (*Adora2a^+^* SPNs: 35 genes; *Drd1^+^* SPNs: 96 genes; **Figure S12D, Table S5,6**).

In addition to this major distinction, the Drop-seq data predict additional eSPN diversity, including an ultra-rare eSPN *Adora2a*^+^ subtype (13-5) that accounts for just 0.3% of all SPNs (n=88 cells). To validate the presence of this ultra-rare type and support the full spectrum of eSPN subcluster diversity implicated by Drop-seq, we performed two triple smFISH experiments for markers of subcluster 13-5. We confirmed instances in the striatum where subcluster 13-5 markers [*Th* and *Npffr1*](http://dropviz.org/?_state_id_=2de1c9ef1e39afed) were co-expressed with each other and with the pan-eSPN marker *Otof* (**Figure 11H**). One clue about the anatomical identity of eSPNs comes from this small Th^+^ population, as spiny Th^+^ principal cells have been observed in striatum with similar spatial arrangement to eSPNs and appear to be dynamically regulated by dopamine (Darmopil et al., 2008).

We conclude that 1) the eSPN vs SPN distinction represents a third axis of SPN diversity, orthogonal to the dSPN/iSPN and patch/matrix SPNs distinctions; 2) eSPNs harbor rare, additional molecular diversity; and 3) by using markers thought to exclusively distinguish iSPNs from dSPNS, functional studies have lumped eSPNs in with canonical SPNs, in spite of their considerable transcriptional divergence. These results highlight the utility of unbiased, high-throughput single-cell methods for defining neuronal populations, even in brain regions and cell populations that have been extensively studied.

## DISCUSSION

The mammalian brain is composed of synaptically interconnected regions, each of which contains a complex mosaic of spatially intermixed cell classes and types. Since Cajal and Golgi, single-cell analyses of cell morphology, cell membranes, and synaptic properties have formed the foundation of our understanding of how structure relates to function in neural circuits. The advent of high-throughput single-cell molecular techniques – such as genome-wide transcript counting with Drop-seq – allow newly systematic approaches to cataloging the essential units of the brain mosaic. Here, we analyzed RNA expression in more than 690K individual cells sampled from nine different regions of the adult mouse brain, encompassing all cell classes. We deployed a novel ICA-based method to disentangle technical effects from endogenous biological signals and highlighted several ways in which such data can identify novel brain cell types, ascertain cell states, and clarify the molecular basis of regionalization across brain circuits and cell classes.

While single-cell profiling provides an exciting opportunity to aid the interrogations of neural circuits, it also presents numerous challenges that we sought to address here. The representations of cells in single-cell datasets can bear an uncertain relationship to their representation in intact complex tissues: we observed quantitative distortions in the representations of cell classes and cell types, driven by library size and cell viability. Such representational issues will be important to consider thoughtfully in future studies in which quantitative cell representation is a key outcome variable, for example, in studies of disease states.

The clusters of cells that are derived from computational analyses cannot be reflexively equated with cell “types”. We identified many patterns of RNA expression that corresponded to cell types, but also many that appear to correspond to cell states or to reflect spatial locations in ways that are continuously varying rather than categorical. The examples described above highlight diverse sources of transcriptional variation among individual cells. Comparisons of expression patterns between cell populations can reflect combinations of these influences, and it will be important to be able to parse these component contributions. We developed an analytical approach to help make signal identification – and its relationship to subclusters – more transparent. This approach allowed us to identify a ubiquitous transcriptional program we believe is enacted to maintain axon and presynaptic function, to different degrees both within and across neuron types. We were also able to resolve signals from striatal SPNs representing differences in pathway (“direct” versus “indirect”), spatial arrangement (“patch” versus “matrix”), and eccentricity—a novel molecular SPN distinction we identified in our dataset. The analytical approach we used was critical for being able to recognize such distinctions and be confident of their biological origin.

The size and complexity of single-cell datasets can limit their utilization by researchers in neuroscience or genetics. To facilitate the utilization of such data throughout neurobiology, we developed an interactive web-based software (DropViz. http://dropviz.org/) that facilitates access and dynamic, responsive exploration of the atlas data.

An exciting direction will be to identify functional and anatomical characteristics that correspond to these transcriptional signatures. We hope that high-volume single-cell gene expression profiles, and the patterns present in such profiles, can function as a *lingua franca* for discussing cellular diversity of the adult mouse brain.

## Footnotes

**Author Contributions** A.S., E.M. and S.A.M. designed the study and wrote the paper. A.S. developed the tissue preparation protocols. A.S., E.M. and M.G. performed Drop-seq and sequenced libraries. A.S, E.M, J.N, N.K., A.W. and S.A.M. developed the ICA analysis pipeline. A.S. and E.M. analyzed the data. D.K. developed DropViz software with help from S.B. M.B. and E.B. performed smFISH experiments. F.K. performed stereological count experiments. S.W. performed the nicotinic receptor analysis. A.G. assisted with cortex regionalization analysis.

## Acknowledgments

This work was supported by the Broad Institute’s Stanley Center for Psychiatric Research and by a Helen Hay Whitney Fellowship to A.S. We thank Rahul Satija and Andrew Butler for advice on single-cell analysis. M.B. was supported by T32GM007753, T32MH020017, HST Idea^A^2, The Sackler Scholar Programme in Psychobiology. We thank Dr. Christina Usher for assistance with manuscript preparation and Dr. Gordon Fishell for comments on manuscript drafts.

